# Transcriptional regulation of *ZIP* genes is independent of local zinc status in Brachypodium shoots upon zinc deficiency and resupply

**DOI:** 10.1101/2020.11.23.393827

**Authors:** Sahand Amini, Borjana Arsova, Sylvie Gobert, Monique Carnol, Bernard Bosman, Patrick Motte, Michelle Watt, Marc Hanikenne

**Author notes:** School of BioSciences, University of Melbourne, Parkville, VIC, 3010, Australia. Corresponding author: Marc Hanikenne.

## Abstract

The biological processes underlying zinc homeostasis are targets for genetic improvement of crops to counter human malnutrition. Detailed phenotyping, ionomic, RNA-Seq analyses and flux measurements with ^67^Zn isotope revealed whole plant molecular events underlying zinc homeostasis upon varying zinc supply and during zinc resupply to starved *Brachypodium distachyon* (Brachypodium) plants. Although both zinc deficiency and excess hindered Brachypodium growth, accumulation of biomass and micronutrients into roots and shoots differed depending on zinc supply. The zinc resupply dynamics involved 1893 zinc-responsive genes. Multiple ZIP transporter genes and dozens of other genes were rapidly and transiently down-regulated in early stages of zinc resupply, suggesting a transient zinc shock, sensed locally in roots. Notably genes with identical regulation were observed in shoots without zinc accumulation, pointing to root-to-shoot signals mediating whole plant responses to zinc resupply. Molecular events uncovered in the grass model Brachypodium are useful for the improvement of staple monocots.

## Introduction

Plants have developed a sophisticated zinc homeostasis network to ensure appropriate zinc supply to tissues throughout their lifetime in varying environments (Choi & Bird, 2014; Clemens et al., 2002). Zinc is an essential micronutrient with catalytic, regulatory and structural functions in enzymes and proteins (Broadley et al., 2007; Gupta et al., 2016). Zinc availability to plants in soils is limited in large areas worldwide (Alloway, 2008), limiting primary productivity and the nutritional quality of agricultural products. Zinc deficiency in plants leads to multiple defects, including lower activity of zinc-binding enzymes, higher reactive oxygen species (ROS) production due in part to reduced ROS-detoxifying copper/zinc superoxide dismutase activity, iron accumulation, membrane and chlorophyll damage, and decrease in photosynthetic performance (Vallee & Falchuk, 1993). These defects macroscopically show during growth and development (Broadley et al., 2007; Sinclair & Krämer, 2012). Zinc toxicity from excess exposure can also occur in plants, mostly in anthropogenically-perturbed areas (Jensen & Pedersen, 2006). The main zinc toxicity symptoms include reduced growth and yield, iron deficiency and hence chlorosis, as well as interference with magnesium, phosphorus and manganese uptake, and reduced root growth and root hair abnormality (Broadley et al., 2007; Fukao et al., 2011).

Many molecular actors for root zinc uptake and its transport to different organs and organelles have been identified (Ricachenevsky et al., 2015; Sinclair & Krämer, 2012). Among them, the Zinc-regulated transporter (ZRT), Iron-regulated transporter (IRT)-like Protein (ZIP) gene family includes 15 and 12 members in Arabidopsis (*Arabidopsis thaliana*, At) and rice (*Oryza sativa*, Os), 10 and 7 of which are up-regulated in response to zinc deficiency, respectively (Assunção et al., 2010; Huang et al., 2020; Kavitha et al., 2015; Krämer et al., 2007; Ramesh et al., 2003; Ricachenevsky et al., 2015; Yang et al., 2009). Although ZIP transporters are widely studied, their specific physiological roles in plant metal homeostasis are not completely understood. Several ZIPs are hypothesized to be responsible for zinc cellular uptake and influx into the cytosol (Colangelo & Guerinot, 2006). However, they have their own specialized functionality and localization, and usually display a broad range of metal substrates. Among the monocotyledonous ZIP transporters involved in zinc, as well as other metal, homeostasis are: rice OsIRT1 (Lee & An, 2009), OsZIP1 (Ramesh et al., 2003), OsZIP4 (Ishimaru et al., 2005), OsZIP5 (Lee et al., 2010), and OsZIP8 (Yang et al., 2009), barley HvZIP7 (Tiong et al., 2014), and maize ZmZIP7 (Li et al., 2016). A number of Heavy Metal ATPases (HMAs) are generally responsible for zinc efflux into the apoplast. The Arabidopsis AtHMA2 and AtHMA4 have an important role in zinc root-to-shoot transport (Hussain et al., 2004). OsHMA2 is apparently the only pump serving this function in rice (Baxter et al., 2003; Takahashi et al., 2012). Metal Tolerance Protein (MTP), Major Facilitator Superfamily (MFS)/Zinc-Induced Facilitator (ZIF), Natural Resistance-Associated Macrophage Protein (NRAMP), Plant Cadmium Resistance (PCR), ATP-Binding Cassette (ABC), Yellow Stripe-Like (YSL), and Vacuolar Iron Transport (VIT), are other transporter families, whose some members are involved in zinc and other metal homeostasis in various dicot and monocot species (Hall & Williams, 2003; Ricachenevsky et al., 2015; Sinclair & Krämer, 2012). Moreover, nicotianamine (NA) is an iron, zinc, copper and manganese chelator involved in intracellular, intercellular and long-distance mobility of these metals in monocots and dicots. NA is synthetized by NA Synthase (NAS) and in graminaceous monocot plants (*i.e.* grasses) exclusively, is the precursor for mugineic acid phytosiderophore (PS) synthesis, which are key for iron acquisition but can also bind zinc (Shojima et al., 1990; Takahashi et al., 1999). The contribution of the 4 *NAS* genes in Arabidopsis was characterized in details (Klatte et al., 2009), and the rice *OsNAS3* gene was demonstrated to be respectively up-regulated and down-regulated by zinc deficiency and excess (Ishimaru et al., 2008; Suzuki et al., 2008), indicating similar functionality.

Sensing and signaling of the zinc status within the plant and in its environment, as well as its integration into a transcriptional regulation of downstream players of the zinc homeostasis network are poorly understood in plants. The Arabidopsis AtbZIP19 and AtbZIP23 are the main known regulators of zinc homeostasis in plants. Both belong to the basic leucine zipper domain-containing (bZIP) TF family and regulate the transcription of *ZIP* and *NAS* genes in response to zinc deficiency in Arabidopsis (Assunção et al., 2010). Close homologs from Barley (HvbZIP56, HvbZIP62, Nazri et al., 2017), wheat (*TabZIPF1* and *TabZIPF4*, Evens et al., 2017) and rice (OsbZIP48 and OsbZIP50, Lilay et al., 2020) could rescue an Arabidopsis *bzip19bzip23* double mutant under zinc deficiency, suggesting a shred function in zinc homeostasis. It is suggested that the Cys/His-rich motif of the AtbZIP19 and AtbZIP23 proteins is involved in zinc sensing via direct zinc binding, which would inactivate these TFs under cellular zinc sufficiency (Assunção et al., 2013; Lilay et al., 2019). In order to discover proteins involved in zinc sensing and signaling in plants, the proteome dynamics upon zinc resupply in zinc-deficient Arabidopsis plants was recently investigated (Arsova et al., 2019). Profiling transcriptome and miRNAome dynamics upon zinc deficiency and zinc re-supply for a few days was also shown to have good potentials to reveal novel zinc-responsive genes and miRNAs in rice (Bandyopadhyay et al., 2017; Zeng et al., 2019).

A large amount of zinc homeostasis research has been carried out on Arabidopsis, and then translated to monocotyledonous crop plants. However, Arabidopsis is not the most suitable model to understand zinc in monocots, principally because grasses and dicots (i) possess divergent developmental and eventually anatomical features, and (ii) employ distinct iron uptake strategies, based on either chelated iron(III) or reduced iron(II) uptake, respectively, resulting in distinct interactions with zinc (Hanikenne et al., in press; Kobayashi & Nishizawa, 2012; Marschner et al., 1986). The latter is indeed very important as evidence indicate the interdependence of zinc and iron homeostasis in Arabidopsis (Arsova et al., 2019; Fukao et al., 2011; Pineau et al., 2012; Scheepers et al., 2020; Shanmugam et al., 2012) and in grasses (Chaney, 1993; Suzuki et al., 2006; Von Wirén et al., 1996). Rice, alternatively, is often used as a model for grasses, but it possesses the unique feature of combining both iron uptake strategies (Ishimaru et al., 2006). Zinc and iron homeostasis in rice is thus not representative of the bulk of other grasses; although zinc and iron cross-homeostasis was also reported in rice (Ishimaru et al., 2008; Kobayashi & Nishizawa, 2012; Ricachenevsky et al., 2011; Saenchai et al., 2016).

In this study, we asked whether novel information on zinc homeostasis in monocots can be obtained by using the grass model *Brachypodium distachyon* (Brachypodium). Being most closely related to wheat and barley among the staple crops, with more similar phenology and root development and anatomy than rice and maize (Watt et al., 2009), the information obtained has potential for easier transfer to valuable crops. Brachypodium combines many attributes of a good model: small and sequenced genome, short life-cycle (3-6 weeks), easily transformable and genetically tractable (Brkljacic et al., 2011; Vogel et al., 2010). Brachypodium was previously proposed as a model for iron and copper homeostasis studies in grasses (Jung et al., 2014; Yordem et al., 2011).

We examined the response of Brachypodium to zinc excess and deficiency, including detailed growth phenotyping. We present commonalities and divergences in zinc homeostasis and its interactions with other metals, iron in particular, between Brachypodium, Arabidopsis and rice. Additionally, ionome and transcriptome dynamics of Brachypodium upon zinc resupply of zinc-deficient plants shed light on striking aspects of zinc homeostasis in this species: a transient down-regulation followed by up-regulation of *ZIP* and 83 additional genes in roots at early time-points (10-30 minutes) upon zinc resupply, assimilated to a local zinc shock response, and a similar response of *ZIP* and 15 other genes in shoots in the absence of zinc accumulation, an indication of rapid root-to-shoot signaling during zinc resupply.

## Materials and Methods

### Plant material, growth conditions and zinc isotopic labeling

*Brachypodium distachyon* Bd21-3 seeds were used (Vogel et al., 2010). De-husked seeds were surface-sterilized by 70% ethanol for 30 seconds, and 50% sodium hypochlorite and 0.1% Triton X-100 for five minutes. After five washing with sterile water, seeds were stratified in sterile water and in the dark for one week at 4°C. Thereafter, seeds were germinated in the dark at room temperature on wet filter paper for three days. Upon germination, seedlings were transplanted in hydroponic trays and control modified Hoagland medium containing 1 μM zinc (ZnSO_4_) and 10 μM Fe(III)-HBED (Scheepers et al., 2020) for one week. Then 10-day-old Brachypodium seedlings were grown in control or treatment conditions in hydroponic media for three weeks. Three static conditions were used: control condition (1.5 μM zinc), deficiency (0 μM zinc), and excess (20 μM zinc). In addition, after three weeks of zinc deficiency, zinc-starved plants were resupplied with 1 μM zinc and then harvested 10 minutes (10 min), or 30 min, or 2 hours (2 h) or 8 h post resupply to capture the dynamic response to a change in zinc supply. Fresh media were replaced each week and last replaced the day before harvest. Harvest took place in a 2h window at day end. The growth conditions were 16 h light per day at 150 μmol m^−2^ s^−1^, 24°C. In all experiments and conditions, hydroponic trays and solution containers were washed prior use with 6N hydrochloric acid to eliminate zinc traces. This procedure was applied in three independent experiments and the replication level of each analysis is detailed in figure legends. In experiment 1, samples were separately collected for (i) root and shoot phenotyping (Fig. 1 to 3 and S1, S10), (ii) ionome profiling (Fig. 4, 5 and S2, S3) and (iii) RNA-Sequencing (Fig. 6–8 and S4-S7, Table 1, Data S1-S5). Root length measurements were performed using the WinRhizo (Regent Instrument Inc., QC Canada) and PaintRhizo (Nagel et al., 2009) tools. Shoot measurements were performed using a LI-3100C Area Meter (LI-COR, NE, USA). Experiment 2 was performed as experiment 1, with the exception that zinc excess was omitted, for independent confirmation of gene expression profiles (Fig. 9). Finally, in experiment 3, static conditions were 0 μM zinc and 1.5 μM zinc as above, and additionally included 1.5 μM of a heavy non-radioactive isotope of zinc (^67^Zn, Isoflex, CA, USA, catalog Nr. 200121-01). Zinc-starved plants were resupplied and labelled with 1 μM ^67^ZnSO_4_, and then harvested at six time-points upon resupply: 10 min, 30 min, 1 h, 2 h, 5 h and 8 h. The isotope-enriched ^67^Zn solution (25 mM) was prepared by dissolving metal ingot in diluted H_2_SO_4_ (Benedicto et al., 2011). In experiment 3, samples were separately collected for (i) isotope concentration analysis and (ii) gene expression profiling by qPCR (Fig. 10 and S8, S9, Data S6).

**Figure 1.**
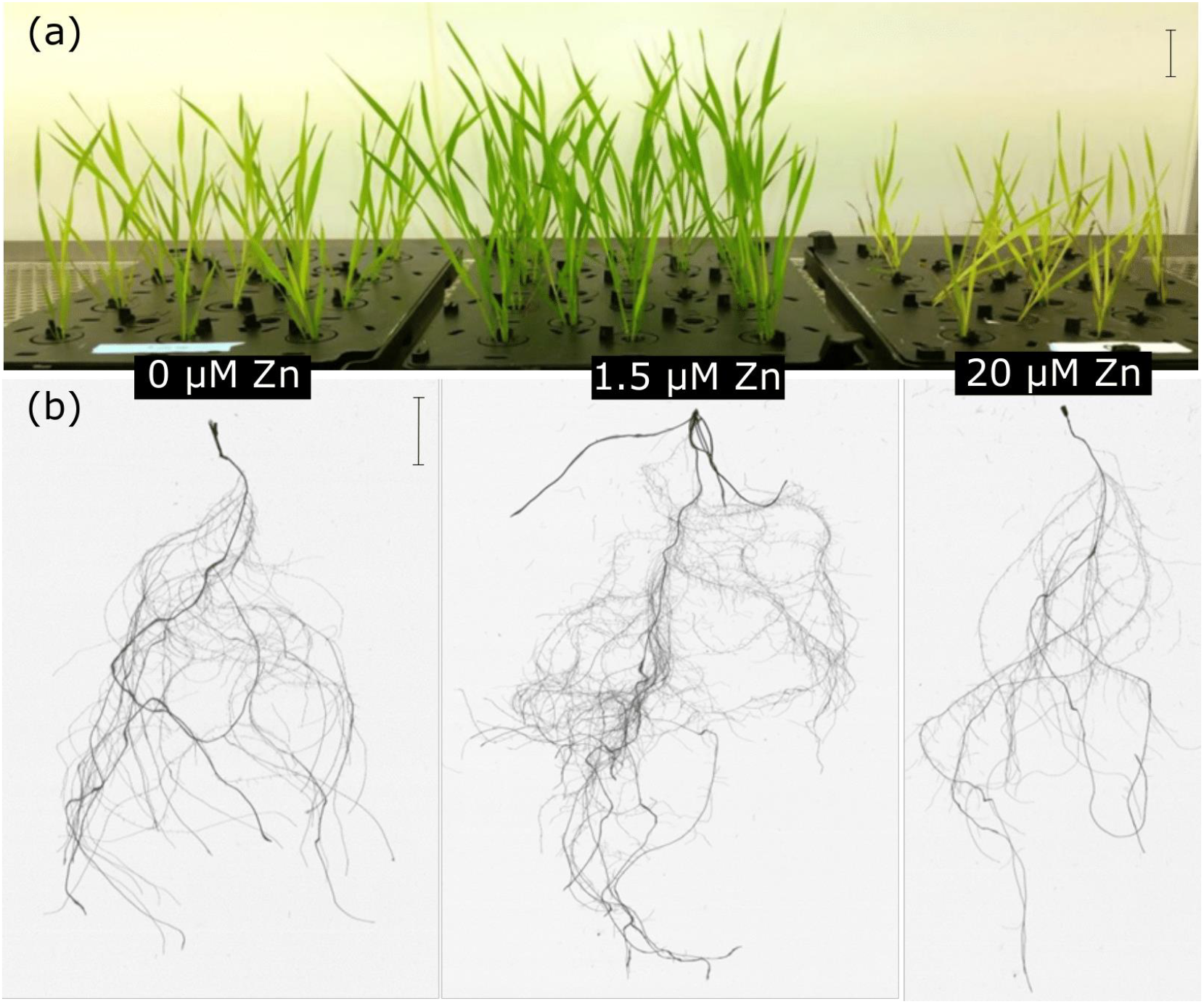
Brachypodium plants grown hydroponically in different zinc regimes for 3 weeks. (a) Shoot and (b) representative root images of plants exposed to zinc deficiency (0 μM Zn, left), control (1.5 μM Zn, center) or excess (20 μM Zn, right) conditions. Pictures are representative of multiple independent experiments. Scale bars are 2 cm.

**Table 1.** Metal homeostasis-related genes among DEG lists

| Brachypodium |  |  |  | Arabidopsis |  | Rice |  |
| --- | --- | --- | --- | --- | --- | --- | --- |
| Gene ID | Description | Tissue | Cluster # | Gene ID | Description | Gene ID | Description |
| Bradi1g53680 | ZIP13 | Root, Shoot | 2, 4 | AT2G32270 | ZIP3 | Os07g12890 | ZIP8 |
| Bradi2g22520 | ZIP5 | Root, Shoot | 2, 4 | AT2G32270 | ZIP3 | Os05g39560 | ZIP5 |
| Bradi2g22530 | ZIP9 | Root, Shoot | 2, 4 | AT2G32270 | ZIP3 | Os05g39540 | ZIP9 |
| Bradi3g17900 | ZIP4 | Root, Shoot | 2, 4 | AT3G12750 | ZIP1 | Os08g10630 | ZIP4 |
| Bradi2g33110 | ZIP7 | Root, Shoot | 2, 4 | AT2G30080 | ZIP6 | Os05g10940 | ZIP7 |
| Bradi1g37667 | ZIP10 | Root, Shoot | 2, 4 | AT1G60960 | IRT3 | Os06g37010 | ZIP10 |
| Bradi1g12860 | IRT1 | Root | 2 | AT4G19690 | IRT1 | Os03g46470 | IRT1 |
| Bradi5g21580 | ZIP3 | Root | 2 | AT3G12750 | ZIP1 | Os04g52310 | ZIP3 |
| Bradi4g26366 | MFS | Root | 5 | AT5G13740 | ZIF1 | Os11g04104 | MFS antiporter |
| Bradi4g43620 | MFS | Shoot | 1 | — | — | Os12g03870 | MFS antiporter |
| Bradi5g08250 | — | Root | 1 | AT5G41000 | YSL4 | Os04g32050 | YSL6 |
| Bradi5g08260 | — | Root | 1 | AT5G41000 | YSL4 | Os04g32050 | YSL6 |
| Bradi1g33347 | HMA1 | Root | 2 | AT4G37270 | HMA1 | Os06g47550 | Cd/Zn-transporting ATPase |
| Bradi1g68950 | Zn/Fe transporter | Root | 3 | AT3G58060 | MTP8 | Os03g12530 | MTP8.1 |
| Bradi4g29720 | — | Root | 2 | AT2G01770 | VIT1 | Os09g23300 | Integral membrane protein |
| Bradi1g17090 | NAS | Root | 8 | AT5G04950 | NAS1 | Os07g48980 | NAS3 |
| Bradi1g53150 | Fe/Mn transporter | Root | 9 | — | — | Os07g15460 | NRAMP6 |
| Bradi5g12456 | PCR11-related | Root | 7 | — | — | — | — |
| Bradi2g43120 | ABC transporter | Root | 9 | AT1G15520 | ABCG40/PDR12 | Os01g42380 | ABCG36/PDR9 |
| Bradi2g31261 | ATX2-related | Root | 3 | — | — | — | — |
| Bradi5g04560 | HIPP26 | Shoot | 8 | AT5G66110 | HIPP27 | Os04g17100 | HIPP42 |
| Bradi3g55480 | ATOX1-related | Root | 4 | — | — | Os02g57350 | — |
| Bradi3g44820 | ATOX1-related | Root | 5 | AT3G56240 | Cu chaperone | — | — |
| Bradi5g12930 | ATOX1-related | Root | 4 | AT4G05030 | Copper transport protein family | — | — |
| Bradi5g25440 | ATOX1-related | Root | 4 | — | — | Os04g57200 | heavy metal transport/detoxification protein |
| Bradi3g27550 | ATOX1-related | Root | 3 | AT2G36950 | Heavy metal transport/detoxification superfamily protein | Os10g30450 | HIPP35 |
| Bradi5g12960 | ATOX1-related | Root | 4 | — | — | Os04g39350 | heavy-metal-associated domain containing protein |

Note that among the 27 Brachypodium genes listed here, only *BdHMA1* and *BdHIPP26* have been attributed names in the Phytozome database (https://phytozome.jgi.doe.gov/pz/portal.html). For The nomenclature proposed in an available ZIP family phylogeny study (Evens *et al.*, 2017) was used where, among the nine ZIP genes present in this list, all received the same numbering as their rice homologs, except for *BdZIP13* which related to *OsZIP8*.

### Ionome profiling

Upon harvest, plant root and shoot material were dried at 50°C for four days and then digested with nitric acid (Nouet et al., 2015). In experiment 1, ionome profiling was performed by inductively coupled plasma atomic emission spectroscopy (ICP-OES, Vista AX, Varian, CA, USA; Nouet et al., 2015). In experiment 3, ^67^Zn, as well as ^66^Zn, barium and vanadium (as negative controls) concentrations were measured by Inductively Coupled Plasma Mass Spectrometry using Dynamic Reaction Cell technology (ICP-MS ELAN DRC II, PerkinElmer Inc., MA, USA) (Benedicto et al., 2011).

### Quantitative RT-PCR

Upon harvest, tissues were snap frozen in liquid nitrogen and stored at −80°C. Total RNAs were extracted from root and shoot samples, cDNA preparation and quantitative RT-PCR were conducted as described (Spielmann et al., 2020). Relative transcript level normalization was performed with the 2^−ΔΔCt^ method using *UBC18* (Bradi4g00660) and *EF1α* (Bradi1g06860) reference genes for normalization (Hong et al., 2008). Primers pairs and their efficiency are provided in Table S1.

### RNA sequencing

42 RNA samples from experiment 1 were used for mRNA-Seq library preparation using the TruSeq Stranded mRNA Library Prep Kit (Illumina, CA, USA). Libraries were multiplexed and single-end 100 nt RNA-Seq was performed on a Novaseq 6000 at the GIGA Center (University of Liege, Belgium) yielding on average ~18 million reads per sample. Raw read sequences were archived at NCBI (Bioproject PRJNA669627). The FastQC software v.0.10.1 (http://www.bioinformatics.babraham.ac.uk/projects/fastqc/) was used for assessing read quality. Trimmomatic tool v.0.32 (Bolger et al., 2014)) was used for removing sequencing adaptors, polyA and low-quality sequences with the following parameters: remove any reads with base with Q < 25 in any sliding window of 10 bases, set crop parameter to 98, leading and trailing to 25, and minimum length to 90 bases. These parameters discarded ~2% of all reads. Using the HISAT2 software v.20.6 (Pertea et al., 2016), reads were mapped on the Brachypodium genome (v.3.2 downloaded from the Phytozome v13 database on 14/02/2020) with --max-intronlen 30000. The average “overall alignment rate” was 93.87% and 98.51% for root and shoot reads, respectively. Finally, mapped read counts (Data S1) were calculated using HTSEQ-COUNT v.0.6.1p1 (Anders et al., 2014).

### Data analysis

The DESEQ2 package v.1.26.0 in R v.3.6.2 (Love et al., 2014) was used for normalizing count data, identification of differentially expressed genes (DEG) with a threshold of absolute fold change of +/−2 and adjusted (Benjamini-Hochberg multiple testing corrected) *p*-value < 0.05, and for principal component analysis (PCA). Gene ontology (GO) enrichment study was performed using the g:GOSt tool embedded in the g:Profiler web server (Raudvere et al., 2019) with threshold of adjusted *p*-value < 0.05, and then visualized with R. For DEG *k*-means clustering, the multiple experiment viewer (MeV) tool was used (Howe et al., 2011). Other statistical tests were conducted using ANOVA or Student’s T-test (see Figure legends).

## Results

### Zinc deficiency and excess hindered Brachypodium shoot growth

Zinc deficiency and excess treatments had negative effects on shoots. Zinc-deficient plants were slightly chlorotic and were shorter than control plants (Fig. 1A), with 22.7% and 27.6% reduction of shoot fresh and dry weight (Fig. 2A-B), as well as 17.5% smaller total leaf area (Fig. 2C). Interestingly, zinc-deficient plants had as median two leaves more than control plants (Fig2D), but with 26.8% lower dry weight per leaf (Fig. S1A).

**Figure 2.**
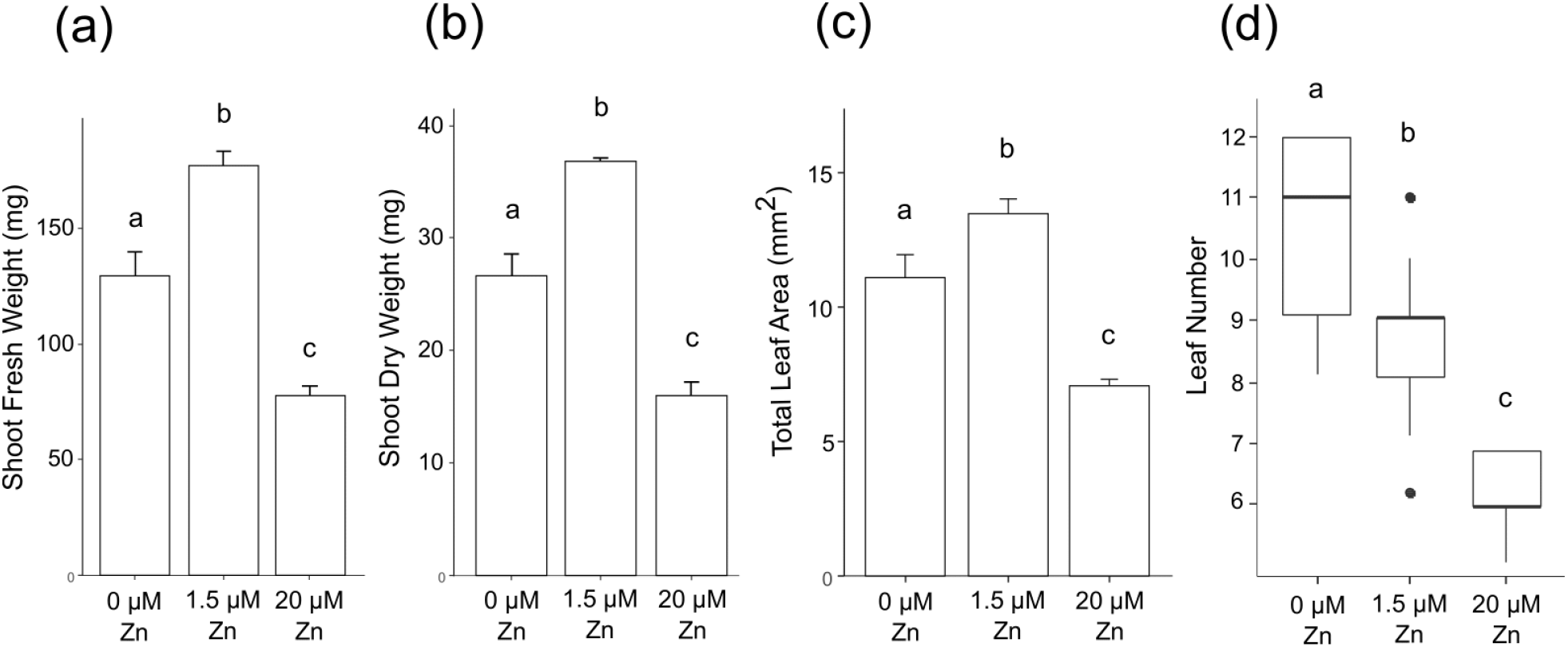
Shoot phenotype of Brachypodium plants upon zinc deficiency and excess. Plants grown hydroponically were exposed for 3 weeks to zinc deficiency (0 μM Zn), control (1.5 μM Zn) or excess (20 μM Zn) conditions. (a) Shoot fresh and (b) dry weight. (c) Total leaf area. (a-c) Bars show mean values (+/− standard deviation) of 9-12 individual plants for each treatment. (d) Leaf number per plant. Box and whisker plot showing the median (hardline), interquartile (box), 1.5 interquartile (whiskers) and outliers (dots) of values from 12 individual plants for each treatment. Letters indicate statistical differences (*p*-value < 0.05) according to Student’s T-test.

Similarly, excess zinc impeded shoot growth and caused leaf chlorosis, but with more severe effects than deficiency (Fig. 1A). Shoot fresh and dry weight as well as total leaf area were 47.7 to 56.2% lower in excess condition compared to control and deficiency (Fig. 2A-C). Plants grown in excess had as median three and five leaves less than control and deficiency plants, respectively (Fig. 2D). Finally, dry weight per leaf of zinc excess plants was 34.3% lower than control plants but the difference with zinc-deficient plants was non-significant (Fig. S1A).

### Zinc deficiency and excess altered root phenotypes of Brachypodium plants

The zinc effect on root growth of Brachypodium plants (Fig. 1B) differed to shoot responses. Under deficiency, root fresh and dry weight, as well as total root length were reduced by 15.2 to 16.5% (Fig. 3A-C). Lateral root length was driving the difference in total root length (Fig. 3D), as it was reduced by 17% while primary root lengths were similar under deficiency and control. Zinc-deficient plants had developed one less nodal root (Fig. 3E) and lower total nodal root length (Fig. S1B) than control plants.

**Figure 3.**
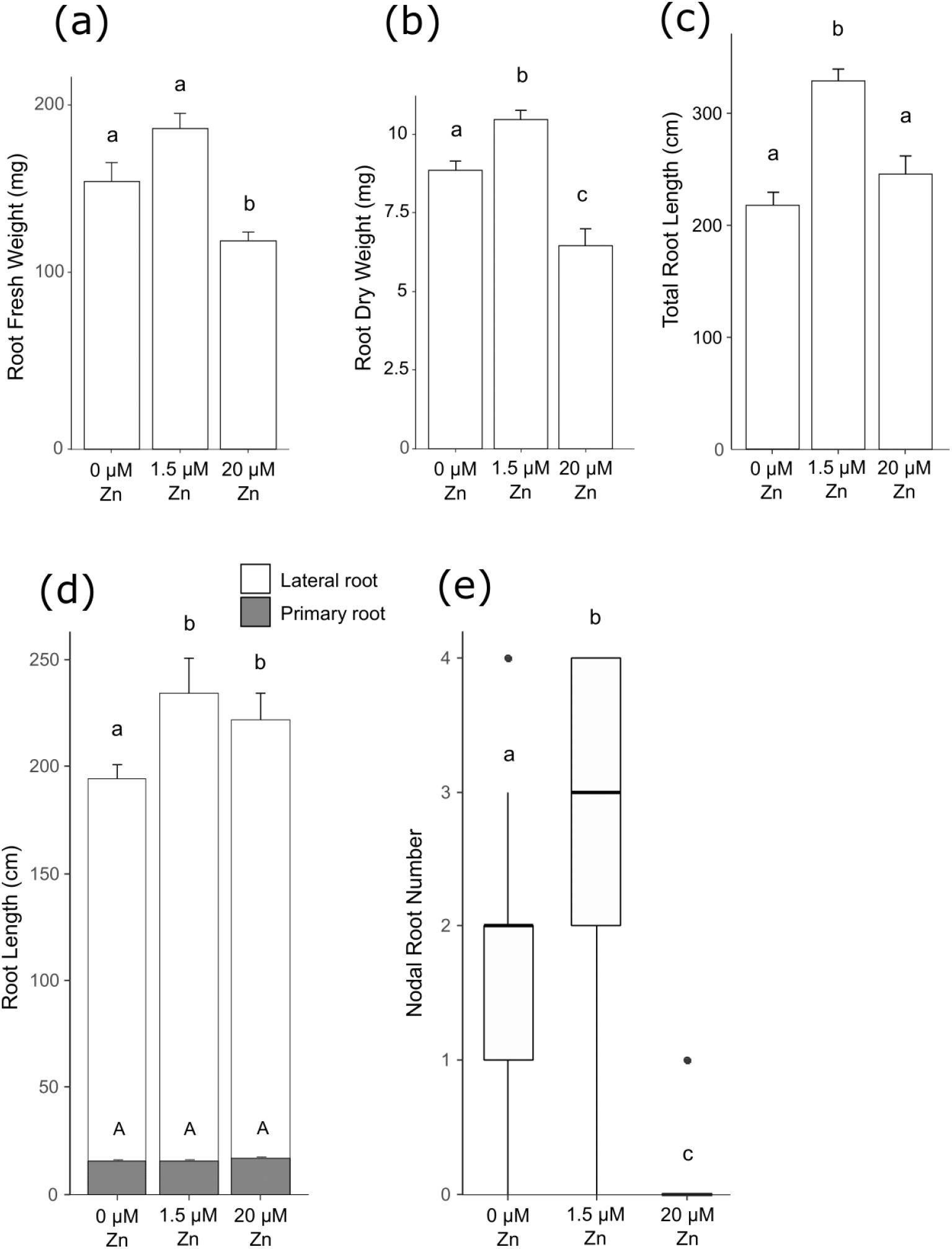
Root phenotypic measures of Brachypodium plants under three weeks of zinc treatments. Plants grown hydroponically were exposed for 3 weeks to zinc deficiency (0 μM Zn), control (1.5 μM Zn) or excess (20 μM Zn) conditions. (a) Root fresh weight, (b) root dry weight, (c) total root length and (d) primary and lateral root length. (a-d) Bars show mean values (+/− standard deviation) of 9-12 individual plants for each treatment. (e) Nodal root number per plant. Box and whisker plot showing the median (hardline), interquartile (box), 1.5 interquartile (whiskers) and outliers (dots) of values from 9 individual plants for each treatment. Letters indicate statistical differences (*p*-value < 0.05) according to Student’s T-test.

Zinc excess plants had 22.3 to 38% lower fresh and dry root weight than control and zinc-deficient plants (Fig. 3A-B), exhibiting stronger responses than the deficiency treatment for these traits. Plants had reduced total root length (Fig. 3C), and slightly (but non-significantly) longer primary roots (Fig. 3D). Total length of lateral roots of zinc excess plants was 5.2% lower than control, but 14.2% higher than in zinc-deficient plants (Fig. 3D). Finally, zinc excess fully inhibited nodal root growth (Fig. 3E and Fig. S1B).

### Zinc deficiency and excess impacted the ionome in Brachypodium

Roots and shoots of zinc-deficient plants had 14.7 and 4.4 times lower zinc concentrations than control plants, respectively (Fig. 4A-B). Zinc excess caused higher zinc accumulation in roots, with 18.6 time greater concentration than in control plants (Fig. 4A). Zinc accumulation in shoots of excess plants was ~4.2 fold higher than in control plants (Fig. 4B). Higher zinc supply corresponded to a greater ratio of root to shoot zinc concentration in Brachypodium, indicating different zinc allocation to the tissues under the different zinc regimes (Fig. 4C).

**Figure 4.**
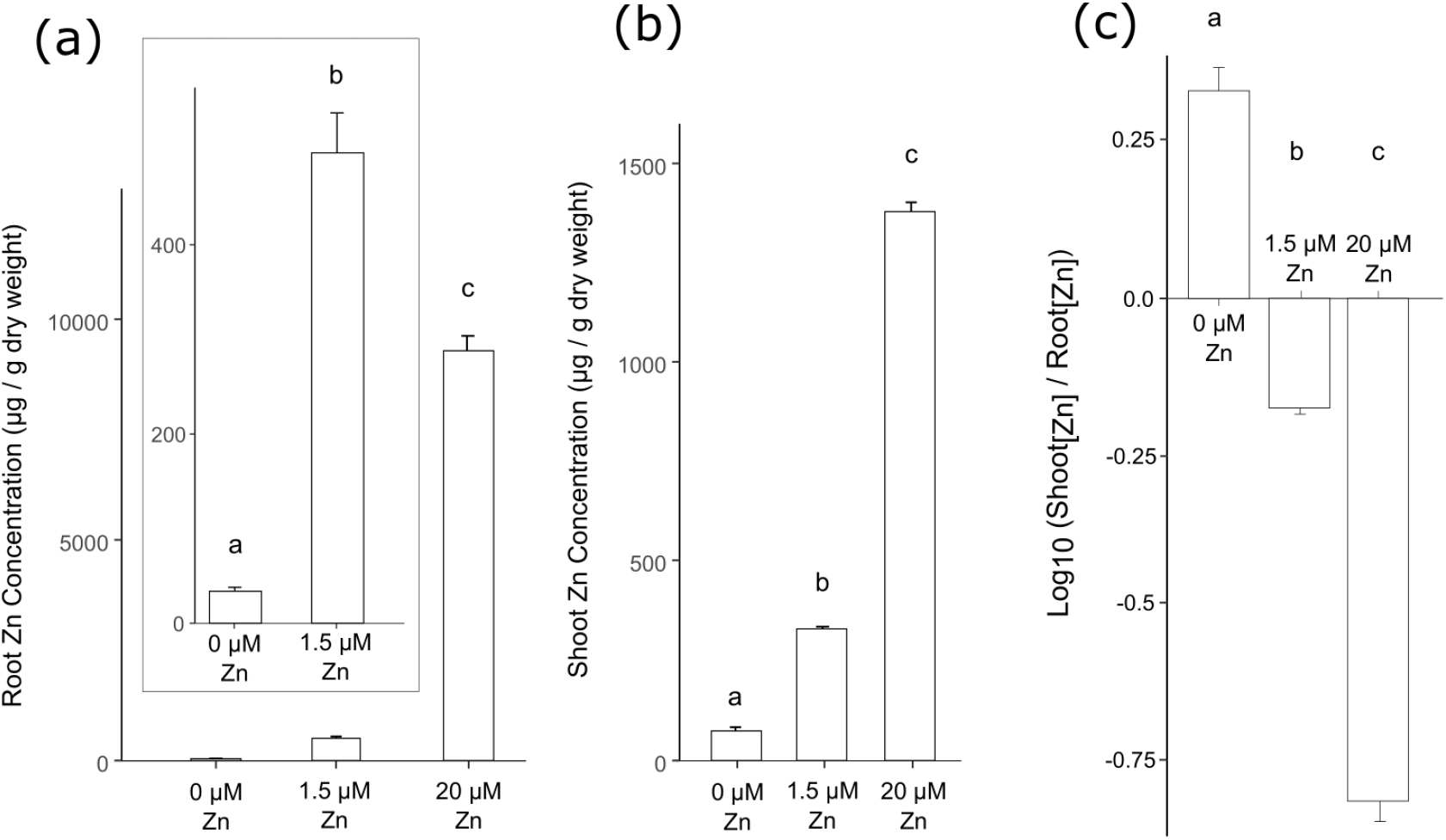
Zinc accumulation in roots and shoots of Brachypodium upon zinc deficiency and excess. Plants grown hydroponically were exposed for 3 weeks to zinc deficiency (0 μM Zn), control (1.5 μM Zn) or excess (20 μM Zn) conditions. (a) Root and (b) shoot zinc concentrations. (c) Shoot to root zinc concentration ratio (Log). Bars show mean values (+/− standard deviation) of three biological replicates (3-4 plants each). Letters indicate statistical differences (*p*-value < 0.05) according to Student’s T-test.

Zinc deficiency and/or excess also affected iron, manganese, copper, calcium and magnesium concentrations in Brachypodium tissues (Fig. S2). Manganese and copper were slightly but significantly less abundant in roots upon deficiency (Fig. S2C,E). Root iron and shoot copper concentrations of zinc-deficient plants were 40.7% increased or 25.3% decreased, respectively (Fig. S2A,F). Notably, upon zinc excess, manganese and copper root concentrations were 3.4 and 2.1-fold reduced compared to control, respectively (Fig. S2C,E). Finally, calcium and magnesium were 33% and 35.1% higher in shoots of zinc excess plants, respectively (Fig. S2H,J).

### Rapid ionome dynamics was observed in Brachypodium roots upon zinc deficiency and resupply

During zinc resupply of zinc deficient roots, we observed gradual accumulation of zinc through time. Zinc increase was modest and non-significant after 10 and 30 min, but reached 3.7-fold after 8 h (Fig. 5A). In contrast to the roots, shoot zinc had no consistent increase within the 8 h of re-supply (Fig. 5A).

**Figure 5.**
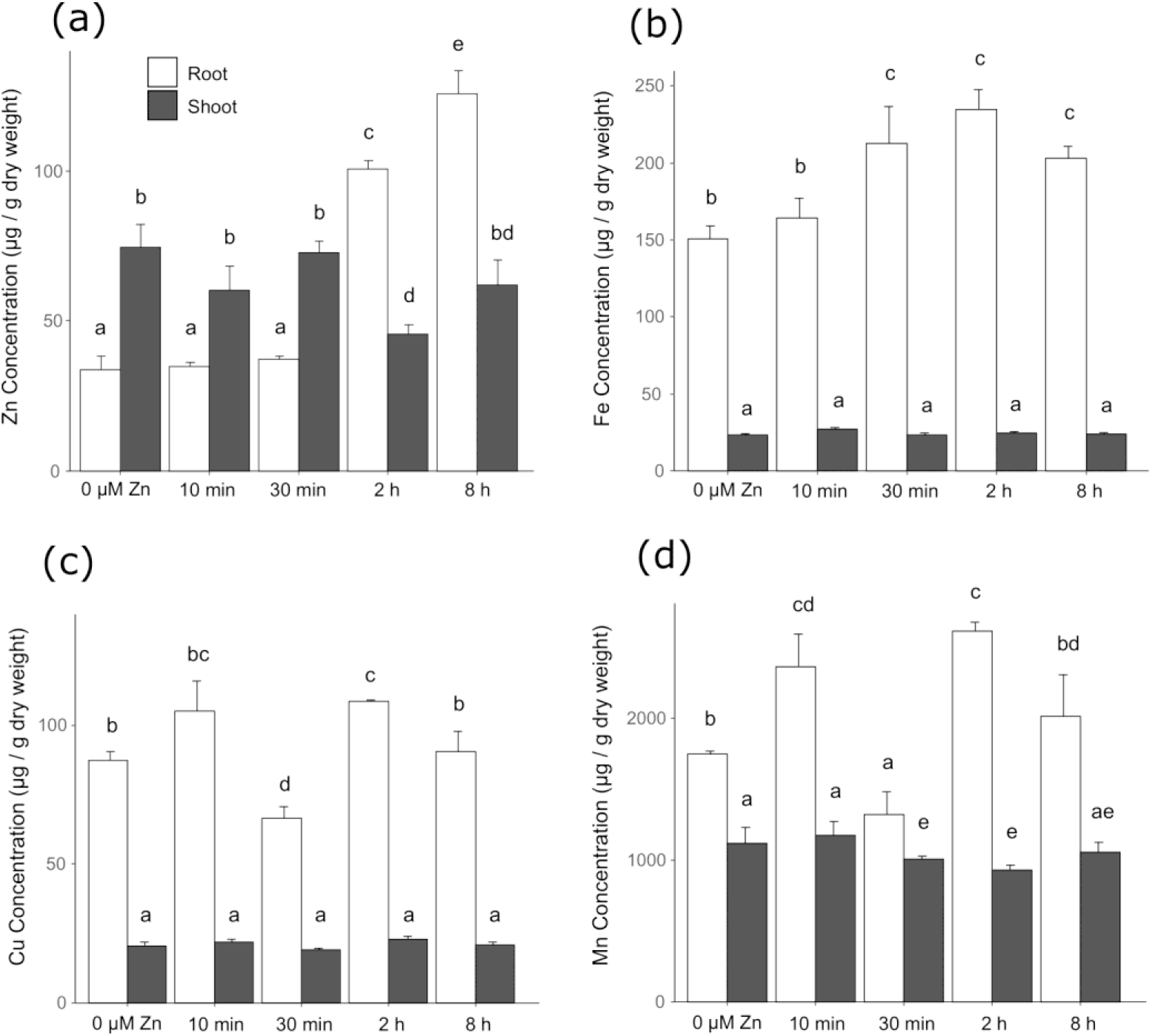
Ionome profiling of roots and shoots of Brachypodium upon zinc deficiency and re-supply. Plants grown hydroponically under zinc deficiency (0 μM Zn) for 3 weeks were resupplied with 1 μM Zn and samples were harvested after short time points (10 minutes to 8 hours). Root and shoot (a) zinc (Zn), (b) iron (Fe), (c) copper (Cu), and (d) manganese (Mn) concentrations. Bars show mean values (+/− standard deviation) of three biological replicates (3-4 plants each). Letters indicate statistical differences (*p*-value < 0.05) according to one-way ANOVA.

Zinc resupply affected the whole ionome (Fig. 5 and Fig. S3). The root iron concentration rose gradually until 2 h parallel to increased zinc level (Fig. 5B). Copper and manganese root concentrations displayed different dynamics to zinc and iron with higher levels after 10 min, but a severe and transient drop at the 30 min time-point (Fig. 5C-D). Changes in the shoot ionome were minor (Fig. 5B-D).

### Transcriptomic responses to steady-state zinc deficiency and excess and to dynamic zinc deficiency and resupply

Principal component analysis (PCA) of the RNA-Seq data indicated that gene expression variance between biological replicates was very low, with the exception of 30 min resupply shoot samples (Fig. 6A-B and S4). In roots (Fig. 6A), samples clustered according to root zinc concentration (Fig. 4, 5). Control and zinc excess samples were similar and zinc deficiency samples distinct (Fig. 6A). Samples collected after 10 min and 30 min of resupply were similar but distinct from deficiency samples. The 2 h and 8 h resupply samples that contained relatively higher zinc concentrations further clustered separately (Fig. 6A). The shoot PCA component(s) affecting sample clustering are more difficult to interpret (Fig. 6B), possibly in relation to the delayed zinc shoot accumulation (Fig. 5A) and therefore lower impact of zinc concentration as a principal component. However, even in shoot, PCA separation between static and resupply samples is clear along PC1, with static conditions on the x-axis left side while resupply samples are progressing with time towards the right along the x-axis (Fig. 6B).

**Figure 6.**
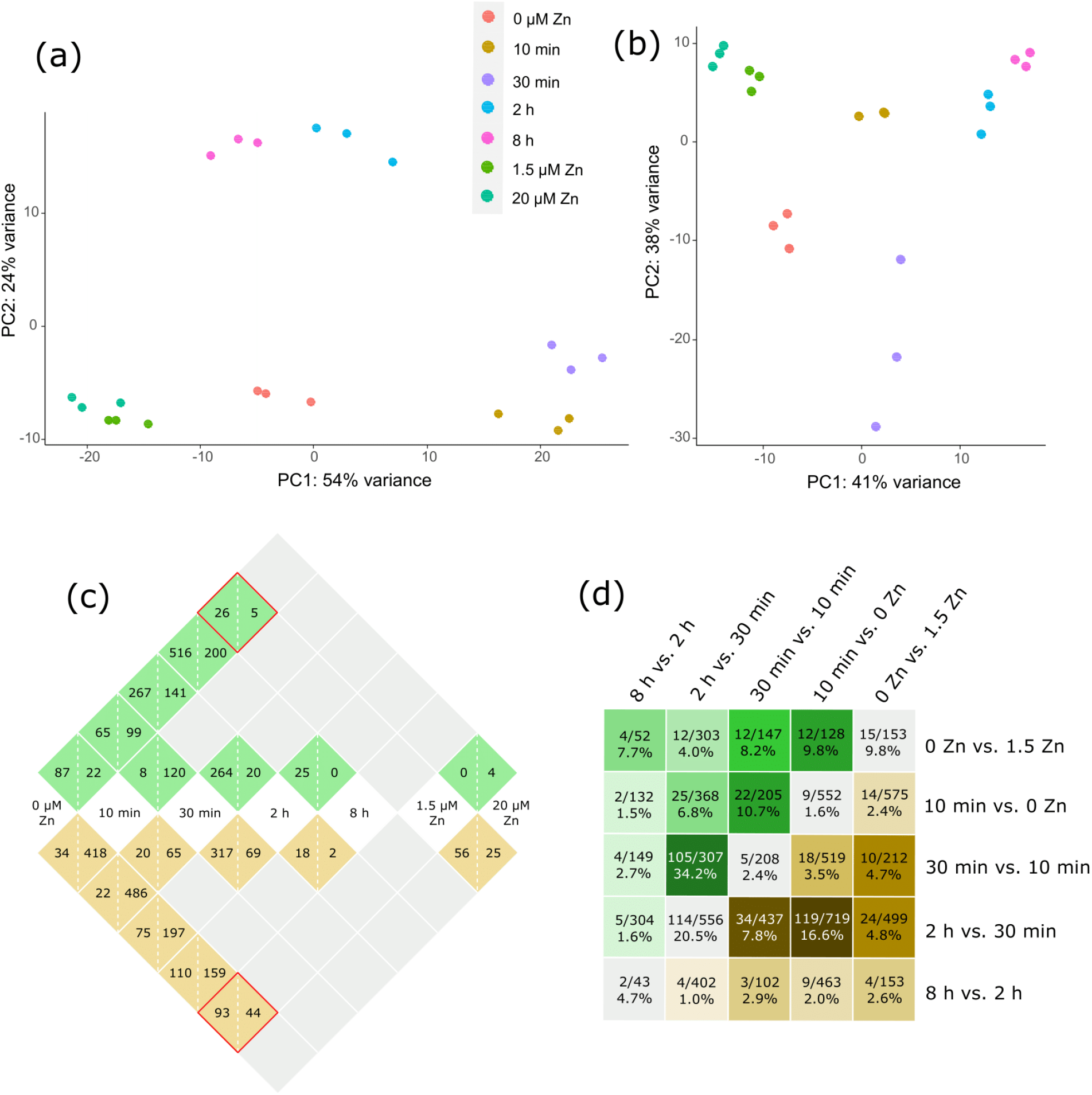
RNA sequencing analysis of the steady-state response to zinc deficiency and excess and the dynamic response to zinc deficiency and resupply in Brachypodium. Data are from three biological replicates (3-4 plants each) for each treatment. Principal Component Analysis (PCA) of (a) root and (b) shoot expression data. PCA of root and shoot data together is presented in Fig. S4. (c) Number of Differentially Expressed Genes (DEG) in 9 selected contrasts in roots (brown cells) and in shoots (green cells). Growth conditions are annotated in central white cells and DEGs identified in a contrast between 2 conditions are annotated in the intersecting cell, with numbers of up-(left) and down-(right) regulated genes. For example, in roots, 93 and 44 genes are respectively up- and down-regulated by zinc deficiency (0 μM zinc) compared to the control condition (1.5 μM zinc) condition (root red square). In shoots, these numbers are respectively 26 and 5 (shoot red square). (d) Ratios of common DEG to the total number of unique DEG for five selected comparisons (deficiency vs. control, and four consecutive comparisons upon zinc resupply) within each tissue are illustrated in green/brown cells. These ratios are also expressed as percentage in each cell. The green upper half of the figure shows shoot data, and the brown lower half shows root data. Color density illustrates the extent of DEG overlap between two comparisons (a darker color corresponding to a larger overlap). The gray diagonal cells present ratios of common DEG to the total number of unique DEG in each comparison between root and shoot tissues.

Differentially expressed genes (DEG) were identified [adjusted *p* < 0.05 and log_2_(fold change) > 1] in a selection of 9 out of 21 possible contrasts between the seven treatments for roots and shoots. The 9 contrasts included comparisons of zinc deficiency and excess to the control (2 comparisons), of zinc resupply time-points to deficiency (4 comparisons), and between consecutive resupply time-points (3 comparisons) (Fig. 6C). 1215 and 976 unique DEG were identified in roots and shoots, respectively. 298 genes were common among roots and shoots, meaning that 1893 unique DEG appeared in 9 contrasts (Data S2). The steady-state responses to zinc deficiency and excess mobilized less DEG than the dynamic response to zinc resupply. There was little overlap in the zinc resupply response between roots and shoots (Fig. 6D). The transcriptional response to zinc resupply was rapid and massive in roots, with the up- or down-regulation of > 450 genes within 10 min (Fig. 6C), with only a small overlap (2.4%) with static deficiency response (Fig. 6D). This latter figure was higher for shoots (9.8%) but differential expression between deficiency and 10 minutes of zinc resupply concerned many genes as well. In roots and shoots, the response to zinc resupply continued to mobilize new genes with time, but slowed down, with a remarkable low number of DEG between the 2- and 8 h time-points (Fig. 6-C-D).

### Unique biological pathways were involved in the dynamic response to Zn resupply compared to steady-state zinc deficiency and excess

DEG, up-regulated and down-regulated, were submitted to Gene Ontology (GO) enrichment analyses. Over-represented biological processes (BPs, *p*-value < 0.05) were identified in most contrasts (Fig. 7, Data S3).

**Figure 7.**
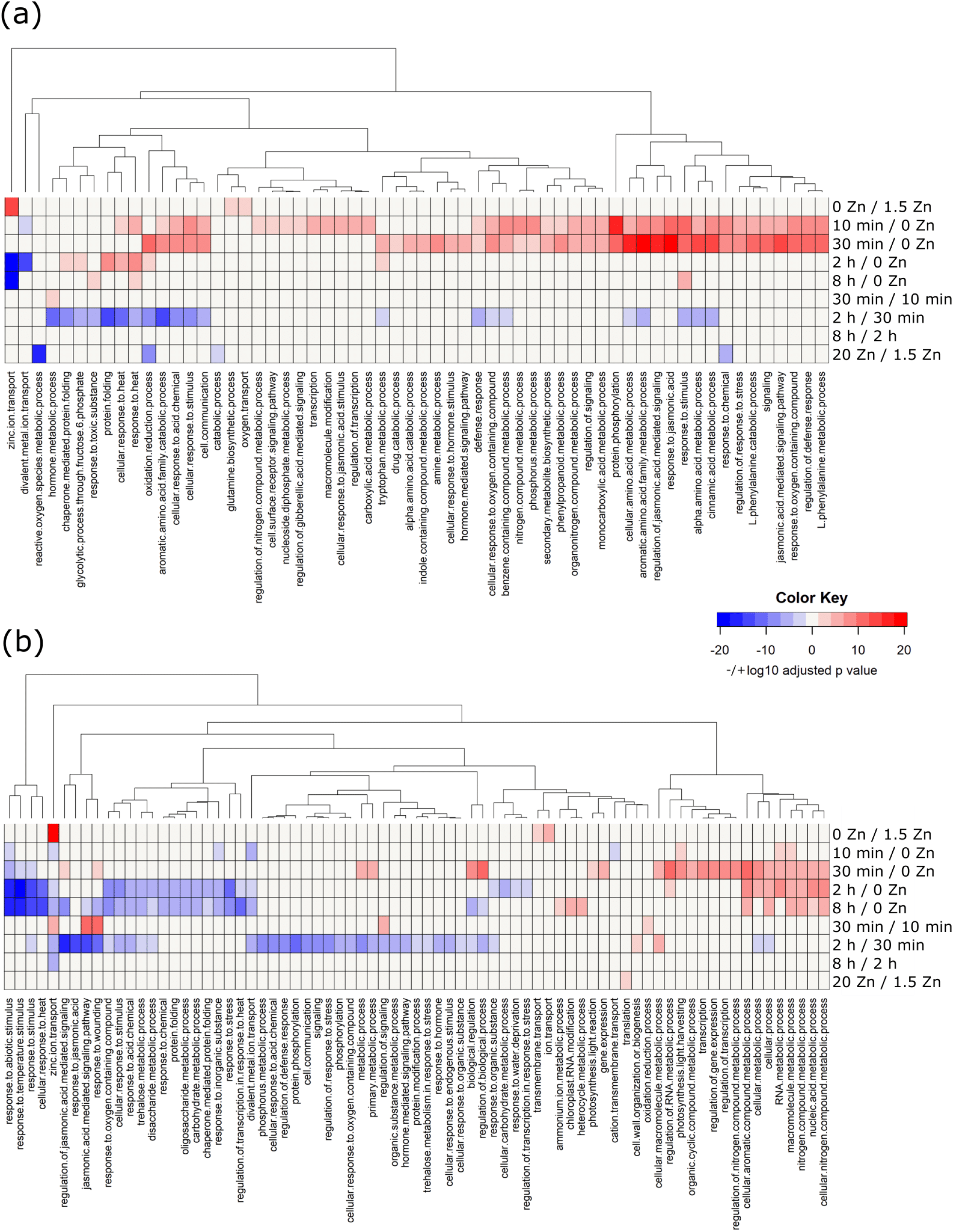
Gene Ontology enrichment analysis of the steady-state response to zinc deficiency and excess and the dynamic response to zinc deficiency and resupply in Brachypodium. The heatmaps present statistically enriched (adj. *p* < 0.05) Biological Processes (BPs) among both up- and down-regulated genes in 9 selected contrasts in roots (a) and in shoots (b). Each row shows a contrast. In the heatmap, the color density indicates the statistical significance of the BP enrichment (-logarithm of adj. p-value), while blue (down) and red (up) colors show the direction of regulation of genes involved in that BP (each BP was specifically only up- or down- regulated with no genes behaving in the opposite direction from that indicated).

The “zinc ion transport” BP was strongly overrepresented among DEGs in roots and shoots of deficient plants (0 vs 1.5 μM zinc), with only up-regulated genes, whereas overrepresentation of catabolism, oxidation-reduction and response to chemical processes was observed in response to excess (20 vs 1.5 μM zinc) in roots, driven by down-regulated genes only (Fig. 7A).

The dynamic response to zinc resupply mobilized many more BPs. In roots, multiple enriched BPs corresponding to up-regulated genes were noticeable (high density red color, Fig. 7A) at the 10 min and/or 30 min time-points compared to deficiency. These BPs were mainly related to signaling, different metabolisms, and stress and hormone responses. “Transcription” as well as other signaling-related BPs were enriched only after 10 min resupply. Noticeably, a single enriched BP, “divalent metal transport”, corresponded to down-regulated genes at 10 min (blue cell at “10 min vs 0 μM zinc”). This item, as well as “zinc ion transport”, was also enriched with down-regulated genes (blue cells) after 2 h of zinc resupply compared to deficiency. As expected, the genes corresponding to the “zinc ion transport” BP were strongly up-regulated at deficiency, but were down-regulated within 2 h upon resupply. Finally, a single or no BP were enriched in “30 vs 10 min” and “8 vs 2 h” consecutive time-point comparisons, respectively, whereas a shift in the zinc resupply response was observed between 30 min and 2 h, with many of the early up-regulated genes being down-regulated in that interval (see the blue cells in the “2 h vs 30 min” comparison, Fig. 7A).

In shoots (Fig. 7B, Data S3), the most striking observation was an enrichment of “zinc ion transport”, “divalent metal ion transport” and “cation transmembrane transport” BPs, corresponding to down-regulated genes within 10 min of resupply, suggesting that a quick transcriptomic regulation of zinc transporter genes preceded zinc re-entry in shoots (Fig. 5A). This response was transient as the enriched “zinc ion transport” BP corresponded to up-regulated genes (30 min) and then down-regulated genes (2 and 8 h) upon resupply compared to deficiency, respectively. Enriched BPs related to transcription, stress and hormone responses, cellular metabolism and regulation, as well as photosynthesis (Fig. 7A, up-regulated BPs, in red), most of which appeared after 10 min resupply in roots, were observed after 30 min in shoots (Fig. 7B), i.e. with one time-point delay.

### Genes encoding members of all zinc transporter families were differentially regulated through time and throughout the different conditions

Among the 1893 identified DEG (Fig. 6), 27 genes were related to zinc/iron/copper/manganese/cadmium homeostasis/resistance, based Phytozome BLAST annotation (Table 1). As Brachypodium is a relatively new model and metal homeostasis studies on this species are scarce, the majority of these annotations were based on sequence or domain similarities with genes/proteins of other species, especially Arabidopsis and rice. Among these 27 genes, 19 genes were differentially expressed in roots only, 2 in shoots only and 6 in both tissues (Table 1). The transcriptional regulation of these genes among the 9 selected contrasts is provided in Fig. S5. The 27 genes belong to families of metal transporters [ZIP (8 genes), MTP (1 gene), HMA (1 gene), NRAMP (1 gene), VIT (1 gene), PCR (1 gene), ABC transporter (1 gene)], metal chelator synthesis [NAS (1 gene)] and transport [MFS/ZIF (2 genes), YSL (2 genes)] or metal chaperones [ATX (1 gene), ATOX1/HIPP (7 genes)]. In general, all known and major zinc transporter families (Ricachenevsky et al., 2015; Sinclair & Krämer, 2012) had thus at least one representative among DEGs. However, at least some of the 27 DEGs (*e.g.* YSL, NRAMP) may be involved not only in zinc, but also in iron, manganese and/or copper homeostasis.

Root and shoot DEG were then clustered according to their expression pattern. Zinc excess was excluded from the analysis as no metal-related genes were regulated in this condition. A total of 9 and 8 clusters were obtained for roots and shoots, respectively (Fig. S6, S7, Data S4). Metal homeostasis-related genes were distributed in several clusters, with distinct expression patterns, including early or late responses as well as transient regulation, mostly in roots. For instance, root cluster #3 (127 genes) contained 3 metal homeostasis genes: an *MTP* and 2 copper metallochaperones (Data S4). These genes displayed increased gene expression upon zinc resupply, especially at 2 and 8 h. The six root *ATOX1*-related copper chaperones were distributed in three clusters (#3, #4, #5) which included genes induced with different kinetics during resupply (Fig. S6). In contrast, *YSL* family genes clustered together with genes whose expression was intermediate at deficiency and high in control but was transiently repressed during resupply (Cluster #1, Fig. S6).

### Gene clustering showed an unusual temporal regulation of*ZIP* genes upon zinc resupply

Among root clusters (Fig. S6), cluster #2 (91 genes) contained all 8 differentially expressed *ZIP* genes identified in root samples (Table 1), as well as two other metal-related genes (*BdHMA1* and a VIT family gene) (RootZIP cluster, Data S5). Similarly, among shoot clusters (Fig. S7), cluster #4 (21 genes) contained all six *ZIP* genes differentially expressed in shoots (ShootZIP cluster, Data S5). Gene expression patterns in root and shoot ZIP clusters (Fig. 8A-B) had a similar shape with two evident peaks of expression at 0 μM zinc and 30 min and a valley at 10 min, resulting in a V-shape consistent with the observation made in the shoot GO enrichment heatmap for the “zinc ion transport” BP.

**Figure 8.**
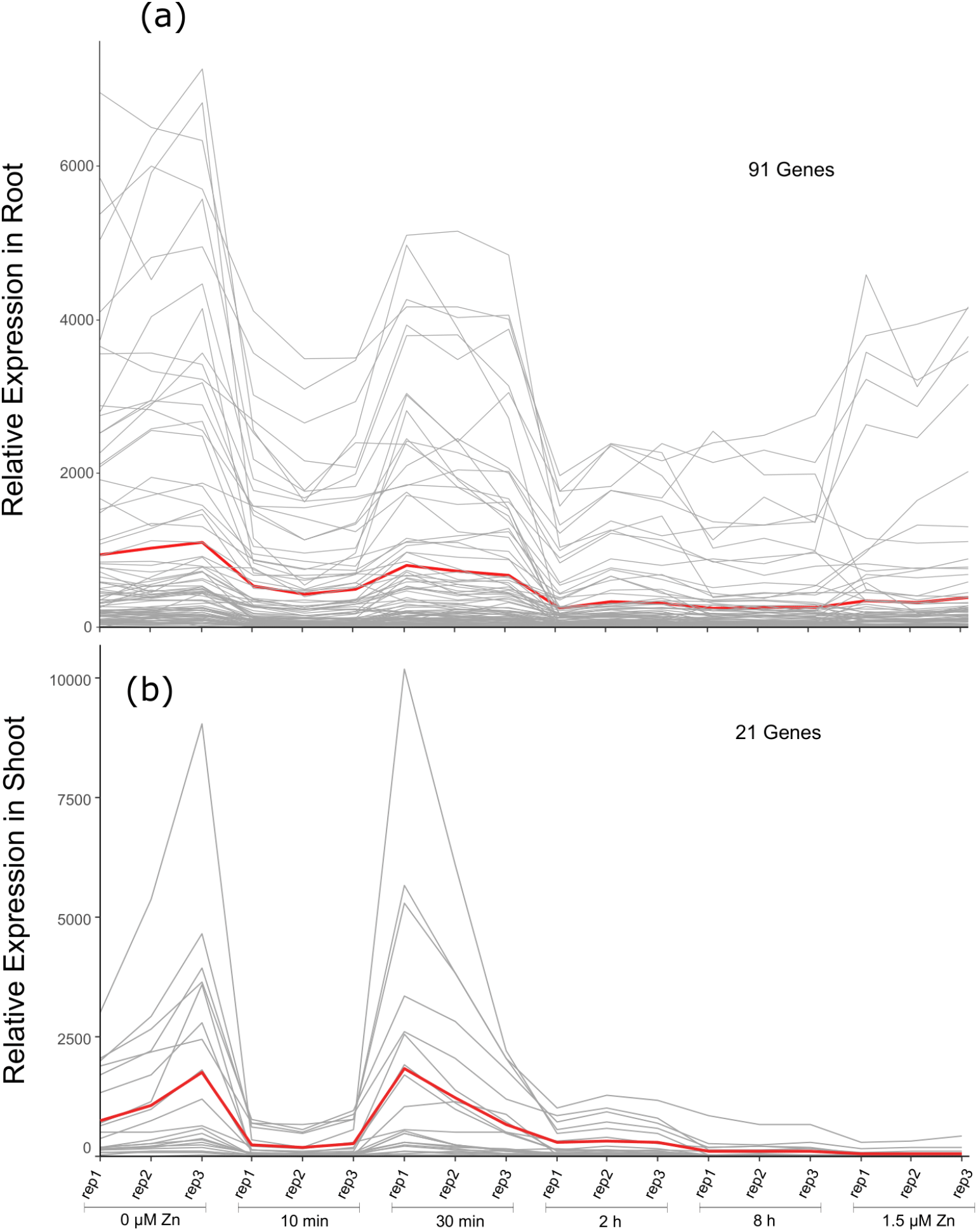
Clustering of gene expression upon zinc deficiency and resupply in Brachypodium. Two clusters containing differentially expressed *ZIP* genes in roots (a) and shoots (b) are shown. Pearson correlation was used as distance metric in *k*-means clustering. The number of differentially expressed genes included in each cluster is noted in each panel. Full clustering data of root and shoot DEGs are shown in Fig. S6 and Fig. S7, respectively. Lines are there to indicate the expression profile of the genes across the three biological replicates, and they should not be considered as time progression. The red lines show the mean expression of all genes in the cluster.

### Shoot *ZIP* transporter genes are down-regulated before measurable amounts of Zn are transported to the shoot

To confirm the V-shape expression pattern of *ZIP* genes (Fig. 8), a fully independent experiment with the same design was conducted, except with the exclusion of zinc excess (Experiment 2, Methods). Quantitative RT-PCR was used to profile expression of selected genes: (i) *BdZIP4*, *BdZIP7* and *BdZIP13* present in RootZIP and ShootZIP clusters, (ii) *BdIRT1* and *BdHMA1* present in the RootZIP cluster only and (iii) the *NAS* gene that was not present in either of these clusters. Complete consistency was observed between RNA sequencing and qPCR data for all six genes in root and shoot tissues in deficiency, resupply and control (Fig. 9).

**Figure 9.**
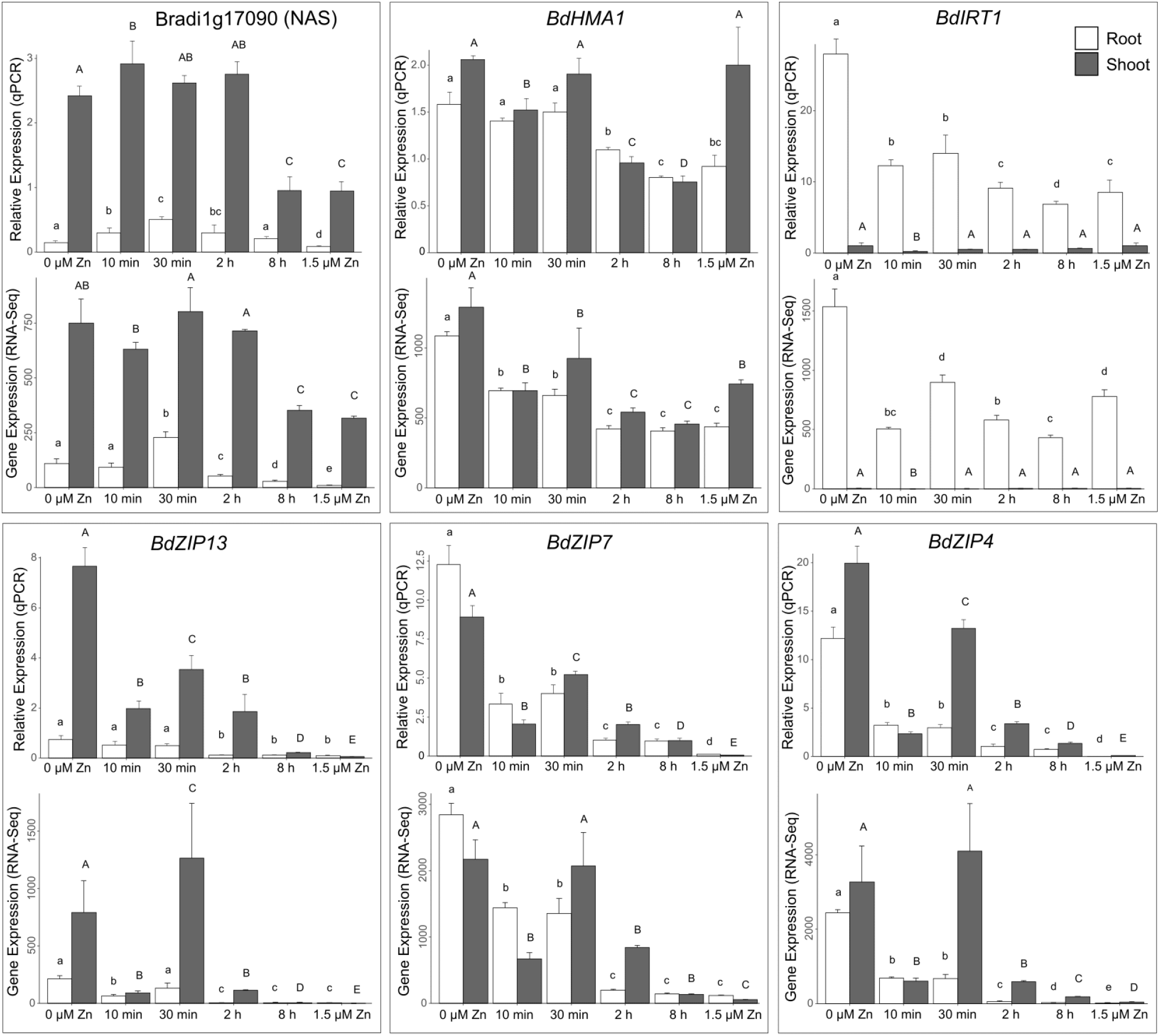
Relative expression of metal homeostasis genes upon zinc deficiency and resupply in Brachypodium. RNA sequencing (RNA-Seq, bottom plot) and quantitative RT-PCR (qPCR, top plot) transcript levels are compared in roots and shoots for the Bradi1g17090 (NAS family), *BdHMA1*, *BdIRT1*, *BdZIP13*, *BdZIP7* and *BdZIP4* genes. Bars show mean values (+/− standard deviation). qPCR expression levels are relative to *UBC18* and *EF1α*, and scaled to average. RNA-Seq and qPCR data that are from two fully independent experiments, each consisting of three biological replicates (2-4 plants each). Letters indicate statistical differences (*p*-value < 0.05) according to one-way ANOVA.

The reproducible V-shape expression pattern of *ZIP* family and 15 other genes in the shoot gene cluster #4 long before zinc influx could be detected in shoots (Fig. 5) was puzzling. A possibility was that a tiny amount of zinc was reaching the shoot tissues rapidly, in an amount lower than the ICP-OES detection limit, and was responsible for local transcriptional regulation for these genes. To enable distinction between zinc still present in shoots after 3 weeks of deficiency (~75 ppm, Fig. 5A) and resupplied zinc, ^67^Zn, a non-radioactive zinc isotope, was used for resupply (Experiment 3, Methods). Note that 1 and 5 h time-points were added to refine the dynamics information. To increase sensitivity and enable detection of zinc isotopes, ^67^Zn concentration measurements were obtained using ICP-MS.

To ensure that ^67^Zn has the same physiological effect as naturally abundant zinc, we first analyzed zinc-responsive genes by qPCR (Fig. S8 compared to Fig. 9). The V-shape expression pattern of *BdZIP4*, *BdZIP7* and *BdZIP13* in roots and shoots were again observed. Second, as natural zinc contains a mixture of stable zinc isotopes, with ^64^Zn being the dominant form and ^67^Zn representing ~4% (Benedicto et al., 2011), natural zinc supply (1.5 μM) was used as a first negative control (Fig. S9A). In line with our expectations, ^67^Zn concentrations were low when plants were treated with natural zinc, and even much lower in deficiency (Fig. S9A). Third, ^66^Zn concentrations in tissues were measured as a second negative control. ^66^Zn measurements were stable throughout the ^67^Zn resupply series whereas it was ~7 times higher when plants were treated with natural zinc (Fig. S9B), as described (Benedicto et al., 2011).

Next, ^67^Zn accumulation in isotope-labelled samples was examined (Fig. 10, Data S6). In roots, a gradual and significant increase of ^67^Zn concentrations was observed with time upon resupply to deficient plants (Fig. 10A, Data S6). The gain in sensitivity compared to Experiment 1 was evident: a significant zinc concentration increase was measured from 10 min (Fig. 10A), when such a change was only detected after 2 h in our initial kinetics (Fig. 5A). In contrast, ^67^Zn accumulation in shoots was only detected after 5 h (Fig. 10B). Examining shoot to root ^67^Zn ratios confirmed that starting from a higher ^67^Zn shoot accumulation in deficiency, ^67^Zn resupply mostly triggered root accumulation up to 5 h before the ratio stabilized (Fig. 10C).

**Figure 10.**
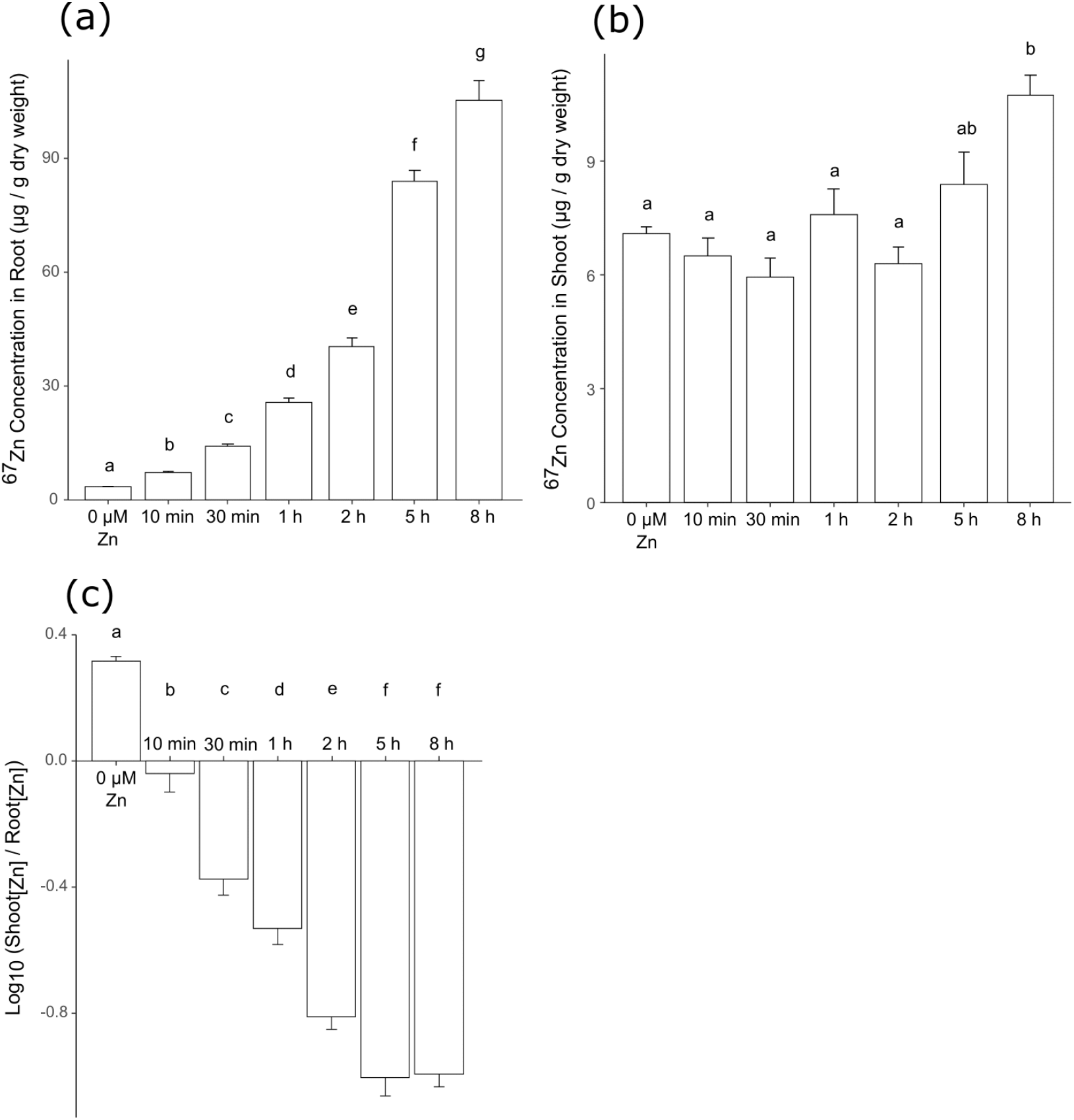
^67^Zn labelling of Brachypodium plants upon zinc deficiency and resupply. Plants grown hydroponically under zinc deficiency (0 μM Zn) for 3 weeks were resupplied with 1 μM ^67^Zn and samples were harvested after short time points (10 minutes to 8 hours). (a) Root, and (b) shoot ^67^Zn concentrations as determined by ICP-MS. (c) Shoot to root ^67^Zn concentration ratio (Log) throughout the time series upon ^67^Zn resupply. Bars show mean values (+/− standard deviation) of three biological replicates (4 plants each). Letters indicate statistical differences (*p*-value < 0.05) according to one-way ANOVA.

## Discussion

In this study, Brachypodium displayed the typical behavior of a zinc-sensitive, excluder plant (Krämer, 2010). It prioritized shoot zinc accumulation upon deficiency and majorly retaining zinc in roots upon excess (Fig. 4), in both cases to preserve the photosynthetic function in leaves. This behavior was very similar to Arabidopsis (Arsova et al., 2019; Talke et al., 2006). However, we showed that the molecular pathways used to achieve this are in part different from Arabidopsis, including distinct interactions (iron) and competition (manganese and copper) with other micronutrients, distinct dynamics of zinc transporter genes and distinct local and systemic signaling.

### Zinc deficiency and excess impact growth and development in Brachypodium

In shoots, increased leaf number was peculiarly associated with reduced total leaf area, total leaf biomass and dry weight per leaf in zinc-deficient plants (Fig. 2B-D and Fig. S1A). Leaf number is known to be influenced by multiple factors such as flowering time and nutrition (Durand et al., 2012; Hu et al., 2017; MacFarlane & Burchett, 2002). In our RNA-Seq data, three homologs of rice flowering-promoting genes, *OsFTL12* (Bradi1g38150, shoots) and *OsFPFL1* (Bradi1g18240, shoots) and *OsFTL6* (Bradi3g48036, roots), were highly up-regulated (7-52 fold) upon zinc deficiency (Data S2). This opens the question of the role of zinc in flowering regulation in Brachypodium. Nutrient deficiency is known to accelerate flowering (Kolář & Seňková, 2008), and flowering is linked with shoot size and leaf number in Arabidopsis, although the effect is variable among early and late flowering ecotypes (Chen & Ludewig, 2018). In Brachypodium, clear repression of vegetative growth was associated with increased leaf number. We hypothesize that in order to optimize nutrient use efficiency in shoot and maintain photosynthesis, plants have adjusted leaf area partitioning (Smith et al., 2017).

Root types were affected differentially depending on zinc supply. Deficiency and excess treatments increased lateral root number and length relative to the primary root, and nodal roots, post-embryonic shoot-born roots emerging from consecutive shoot nodes and a unique feature of monocots, were strongly affected. Their initiation was fully inhibited upon zinc excess (Fig. 3E). Nodal roots of wheat (Tennant, 1976) are strongly suppressed by low nutrients. In Brachypodium, deprivation of nitrogen, phosphorus (Poiré et al., 2014) and water (Chochois et al., 2015) similarly results in significantly lower number of nodal roots. A positive correlation between nodal root numbers and the nutrient uptake, including nitrogen, phosphorus, iron and zinc, is observed in rice (Subedi et al., 2019). Due to a higher diameter of metaxylem to seminal roots and consequent impact on nutrient uptake capacity, nodal roots play a role in nitrate supply to the plant (Liu et al., 2020; Steffens & Rasmussen, 2016). If this is true for zinc too, the observed absence of nodal roots during zinc toxicity can be interpreted as a protective mechanism that minimizes zinc uptake into the plant. However, the decreased number of nodal roots during deficiency does not fully fit into this narrative, unless the development of nodal roots itself has specific zinc requirements. Furthermore, nodal roots provide mechanical stability to the plant (e.g. from winds, Liu et al., 2020), the decreased number or absence of nodal roots in soils with unfavorable zinc conditions may prove to be disadvantageous to logging in various crops and thus further increase of yield loss (in addition to the physiological zinc effects). It would therefpre be interesting to look for variation in nodal root allocation in response to zinc among Brachypodium accessions, as was found for water supply (Chochois et al., 2015).

### Interaction of zinc and other metal homeostasis

Zinc excess had no impact on iron root and shoot levels in Brachypodium (Fig. S2A) and no enrichment for iron homeostasis genes was observed in the transcriptomic response to zinc excess (Fig. 7, Data S2). This contrasts with results from Arabidopsis where zinc excess triggers a secondary iron deficiency with a strong transcriptional response, and zinc toxicity symptoms can be alleviated by higher iron supply (Fukao et al., 2011; Hanikenne et al., in press; Lešková et al., 2017; Shanmugam et al., 2012; Zargar et al., 2015). The iron accumulation dynamics in Brachypodium roots was also in contrast to Arabidopsis with a transient increase upon zinc resupply (Fig. 5B). Zinc deficiency and resupply instead induces a transient decrease in iron concentration in roots of Arabidopsis (Arsova et al., 2019).

Differences to Arabidopsis studies may be because dicot plants and grasses use distinct iron uptake systems (Kobayashi et al., 2012; Hanikenne et al., in press). In dicot plants such as Arabidopsis, iron uptake is based on a reduction strategy where iron(II) is taken-up by IRT1, whereas in grasses, it is based on iron(III) chelation by phytosiderophores (PS) in the rhizosphere prior PS-iron(III) uptake by roots (Hanikenne et al., in press; Kobayashi & Nishizawa, 2012). The chelation strategy provides higher uptake specificity and possibly enables less interference by divalent cations such as zinc, although PS were shown to bind zinc in the rhizosphere (Suzuki et al., 2006). None of the genes involved in the chelation strategy in grasses were among zinc-regulated genes in Brachypodium (Data S2). IRT1 homologs are also found in grasses (Evens et al., 2017) and were shown to transport zinc and iron (Ishimaru et al., 2006; Lee & An, 2009; Li et al., 2015). In this study, and in contrast to *OsIRT1* (Ishimaru et al., 2008), *BdIRT1* was regulated by zinc availability (Fig. S6). With other ZIPs sharing a similar expression pattern, *BdIRT1* may be involved in iron and zinc transport, and be responsible for higher accumulation of iron upon zinc deficiency (Fig. S2A), as well as for the parallel increase of zinc and iron uptake at early time-points upon zinc resupply (Fig. 5A-B).

Competition in root uptake between zinc and manganese/copper was possibly regulated by the same (or another set of) ZIP transporters (Fig. S2C,E). In rice and wheat, similar competition was reported for manganese (Evens et al., 2017; Ishimaru et al., 2008). ZIP, as well as MTP, proteins can indeed potentially transport zinc and manganese (Milner et al., 2013). *AtMTP8* and *OsMTP8.1*, although responding to zinc deficiency, are manganese transporters (Chen et al., 2013; Chu et al., 2017). *BdHMA1*, homolog of *AtHMA1* (Kim et al., 2009; Seigneurin-Berny et al., 2006), a RootZIP cluster gene (Fig. 8), may mediate zinc/copper interactions. Moreover, among the metal homeostasis genes regulated by zinc in Brachypodium (Table 1, Fig. S5), seven are reported to encode proteins related to the human ATOX1 metallochaperone involved in copper chelation (Walker et al., 2002). Annotated as heavy-metal-associated domain (HMAD) containing proteins, these proteins are also known as heavy metal associated isoprenylated plant protein (*HIPP*) genes in plants (de Abreu-Neto et al., 2013). Arabidopsis and rice HIPP homologs were found to be cadmium-responsive and/or involved in copper transport (Shin et al., 2012; Zhang et al., 2018). Representing almost a quarter of metal homeostasis DEGs in our dataset (7/27), ATOX1-related copper chaperones may also be involved in zinc chelation in Brachypodium, indicating a complex metal interplay.

### Transcriptional regulation of the Brachypodium zinc response

The AtbZIP19 and AtbZIP23 transcription factors from Arabidopsis are the best studied regulation system coordinating the zinc deficiency response in plants (Assunção et al., 2010). Homologs with conserved functions were characterized in barley, wheat and rice (Castro et al., 2017; Evens et al., 2017; Lilay et al., 2020; Nazri et al., 2017). The Brachypodium homolog of *AtbZIP19*, Bradi1g30140 [annotated as *BdbZIP9* in Phytozome v.12.1, but as *BdbZIP10* or *BdbZIP11* in (Glover-Cutter et al., 2014) or (Evens et al., 2017)] was previously suggested to be involved in a zinc deficiency-induced oxidative stress response (Glover-Cutter et al., 2014; Martin et al., 2018). However, here, *BdbZIP9* was barely regulated by zinc supply: it was slightly more expressed in zinc-deficient shoots compared to control plants and displayed a very flattened V-shape dynamics upon zinc resupply (Fig. S10A). AtbZIP19 and AtbZIP23 are proposed to be specialized in either roots or shoots, respectively (Arsova et al., 2019; Sinclair et al., 2018). *BdbZIP9* was more expressed in shoots than roots (Fig. S10A). Interestingly, another *bZIP* gene, Bradi1g29920 (*BdbZIP8* in Phytozome v.12.1), was majorly expressed in roots (Fig. S10B) and, although it was not present among initially identified DEG (1.9-fold down-regulation 10 min after resupply, Data S1), it displayed the same V-shape expression pattern as ZIP cluster genes upon zinc resupply, suggesting that BdbZIP8 may be involved in zinc homeostasis in Brachypodium.

Additionally, 113 TFs from various families such as WRKY (25 genes), AP2 (24 genes), MYB (22 genes), bHLH (11 genes), and bZIP (9 genes) were among identified DEG (Data S7). None of them are homologs of known zinc regulatory genes and constitute new candidates for a role in zinc homeostasis regulation in grasses.

### Zinc translocation to the shoot is a slow process

Zinc translocation to shoots was delayed relative to the rapid zinc re-entry in root tissues upon zinc resupply in Brachypodium, similar to earlier observations in Arabidopsis (Arsova et al., 2019). The Arabidopsis AtHMA2 and AtHMA4 pumps, as well as their rice homolog OsHMA2 were shown to be mostly responsible for root-to-shoot zinc transfer (Hussain et al., 2004; Satoh-Nagasawa et al., 2012). Whereas *AtHMA2* expression is induced by zinc deficiency (Arsova et al., 2019; Sinclair et al., 2018), *AtHMA4* and *OsHMA2* expression is barely regulated by zinc (Talke et al., 2006; Wintz et al., 2003; Yamada et al., 2013). Their Brachypodium homolog (Bradi1g34140) was moderately induced by zinc deficiency (1.6 fold) in roots, then transiently down-regulated upon zinc resupply before peaking at after 8 h (Fig. S10C). This up-regulation may be responsible for zinc re-entry observed in shoots after 5 h of resupply (Fig. 10), based on modelling showing that small variations in *HMA4* expression in Arabidopsis suffice to produce large effects in zinc efflux of symplast and to vasculature (Claus et al., 2013). The Bradi1g34140 late induction upon zinc resupply may therefore be responsible for delayed zinc accumulation in shoots. Moreover, the *PCR11*-related gene, found among zinc-responsive genes in Brachypodium (Table 1) was up-regulated at the 10 min and 30 min resupply time-points and gradually down-regulated thereafter (Fig. S6, Cluster #7). It may serve as a minimal shoot zinc supplier when the HMA pump is down-regulated (Song et al., 2010).

In contrast to zinc, copper and manganese concentrations changed quickly upon zinc resupply. Both metals experienced an increase at 10 min and then a decrease at 30 min, the inverse of the V-shape of ZIP clusters in root and shoot, although it was only significant in root (Fig. 5C-D). *OsNRAMP5* is suggested to function in manganese distribution from root into shoot (Yang et al., 2014). The zinc-responsive *NRAMP* gene (homolog of *OsNRAMP6, Bradi1g53150*) may serve the same function in Brachypodium. Its severe induction at 10 min time-point and with excess zinc, where manganese concentration is lowered (Fig. S10D) can support its role in manganese root-to-shoot translocation. On the other hand, *OsATX1*, homolog of *ATOX1*-related copper chaperone, was reported to have an important role in root-to-shoot copper translocation (Zhang et al., 2018) and to interact with multiple rice HMA pumps, probably to transfer copper to these pumps. There are seven *ATOX1*-related genes in the metal list, some of which were immediately regulated by zinc resupply (Clusters #4 in Fig. S6, cluster #8 in Fig. S7). Rapid induction of the *NRAMP* gene and several *ATOX1*-related genes (Fig. S5), in contrast to the late induction of *AtHMA4* homolog (Bradi1g34140), might explain the efficient regulation of manganese and copper concentration in shoot, compared to zinc.

### Zinc shock appears to be the first transcriptomics response upon Zn resupply to deficient roots

Expression patterns of the root ZIP cluster genes (Fig. 8 and 9) were in stark contrast to observations made in Arabidopsis. In Brachypodium, genes within this cluster were highly expressed at zinc deficiency, rapidly down-regulated after 10 min resupply, then up again after 30 min, thus displaying a V-shape expression pattern (Fig. 8 and 9). This response occurred in roots as zinc concentration was steadily increasing upon resupply (Fig. 5A and 10A). In Arabidopsis was observed an initial up-regulation in roots of multiple metal homeostasis genes and proteins, including ZIPs, after 10 min of resupply of zinc-starved plant before a down-regulation from 30 min (Arsova et al., 2019).

The V-shape expression pattern of the ZIP cluster genes in roots implies that zinc influx into roots of zinc-starved plants is sensed as a zinc stress, similar to a zinc excess. This sensing then initiates within 10 minutes down-regulation of zinc uptake genes in roots. Such zinc shock response was described in the yeast *Saccharomyces cerevisiae* (MacDiarmid et al., 2003; Simm et al., 2007). Thereafter, upon sensing yet below-sufficient zinc levels in root tissues, *ZIP* genes are re-up-regulated at 30 minutes followed by more classical down-regulation with increasing zinc concentrations in tissues at later time-points. The response to zinc resupply in roots therefore occurs in two phases (Fig. 6A,D), an initial and rapid phase (10-30 minutes) combining zinc shock response as well as zinc reuptake supported by intense signaling (Fig. 7A), and a later phase (2-8 hours) which corresponds to a slow return to a sufficient state. Although they display very different dynamics, two phases are also observed in response to zinc resupply in Arabidopsis (Arsova et al., 2019).

### Early transcriptomic response of zinc transporter genes in shoots mirrors the root pattern and is independent of local zinc concentration

Strikingly, shoot ZIP cluster genes (Fig. 8 and 9) displayed a V-shape expression pattern as in roots (Fig. 8 and 10) although no change in shoot zinc level can be detected within this time-frame (Fig. 5A and 10B). In Arabidopsis no regulation of ZIPs at transcriptional or translational level was observed in shoots before 8 h of zinc resupply (Arsova et al., 2019). Thus, early transcriptomic response of zinc transporter genes in shoots appears to be independent of local zinc concentration and to be coordinated with roots in Brachypodium and we propose that zinc re-entry in roots initiates a root-to-shoot signaling that instigates a distant transcriptomic response (Fig. 11).

**Figure 11.**
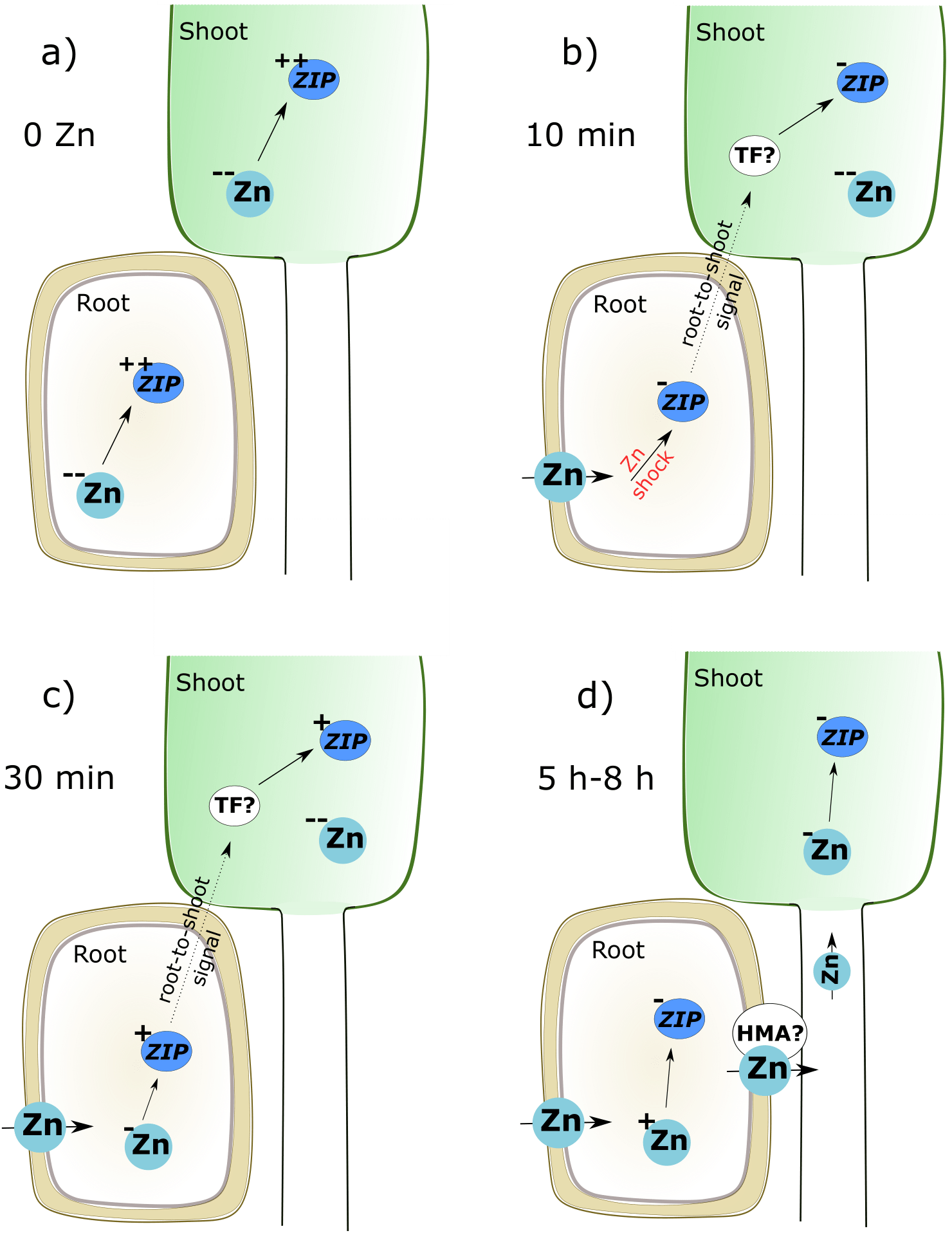
Working model of root-to-shoot signaling upon zinc deficiency and resupply in Brachypodium. (a) Zinc deficiency (0 Zn). Depletion of zinc in root and shoot causes strong upregulation of *ZIP* genes in both tissues. (b) 10 minutes after zinc resupply (10 min). After a depletion period, zinc resupply is sensed as stress (Zn shock) in roots which triggers rapid down-regulation of *ZIP* gene expression in roots and initiates root-to-shoot signaling. In shoots, *ZIP* genes are also immediately downregulated although zinc is not transported to shoot yet. (c) 30 minutes after zinc resupply (30 min). Zinc continues to accumulate in root cells, but remains at low concentration. *ZIP* genes are upregulated again to sustain zinc uptake. This status is signaled to the shoot to induce a similar response. (d) Five to eight hours after zinc resupply (5 h-8 h). Zinc concentration keeps increasing which results in the downregulation of *ZIP* genes. At the same time, zinc is translocated to shoot (probably by an HMA homolog; Bradi1g34140) and accumulation of zinc in shoot cells downregulates *ZIP* genes in shoot as well though local signaling. Double-plus (++) shows very high quantity, plus (+) shows moderate quantity, minus (−) shows low quantity and double-minus (−−) shows very low quantity.

Shoot transcriptome response, independent from shoot nutrient concentration, was reported upon nitrogen resupply to nitrogen-starved maize plants (Takei et al., 2002). In roots, several signaling-related BPs were enriched at 10 min and 30 min time-points (Fig. 7A, dense red area, up-regulated genes), while this response was delayed in shoots where metal transport (Fig. 7B, in blue, down-regulated genes) was among the few enriched BPs after 10 min (Fig. 7B). It is therefore tempting to speculate that the root-to-shoot signaling directly represses expression of metal transporter genes in shoot, rather than activating local signaling pathways in shoot. Supporting this idea is that the “RNA metabolism” BP was also enriched (Fig. 7B, in red, up-regulated genes), 10 min after resupply in shoots. Several transcription factors were found in this enriched BP, and belong to different superfamilies such as B3, AP2, WRKY and bZIP (Bradi4g02570, Data S3). These TFs may potentially regulate *ZIP* genes in Brachypodium shoots upon zinc resupply.

Long-distance signaling mechanisms known in plants include electric or hydraulic signaling, calcium waves propagated by calcium-dependent protein kinases and calmoduline proteins, ROS waves, sugar signaling, hormonal signaling and mobile mRNA (Shabala et al., 2016). Among the signaling-related DEG and enriched BPs at 10 min in root, multiple genes connected to these processes are present and constitute candidates for producing root-to-shoot signals (Data S5).

Long distance or systemic signaling is known to contribute to metal homeostasis regulation. It was suggested that *AtMTP2* and *AtHMA2* transcript levels in roots are regulated by shoot zinc concentration in Arabidopsis, in contrast to *ZIP* genes being controlled by the local zinc status (Sinclair et al., 2018). Designing an experiment testing our model of root-to-shoot signaling upon nutrient resupply is a challenge. Split-root experiments were successful to disentangle local versus systemic signals regulating the response to iron deficiency (Schikora & Schmidt, 2001; Vert et al., 2003; Wang et al., 2007). To study systemic shoot-to-root signaling, reciprocal grafting of mutant and wild-type roots and shoots, and foliar nutrient supply are popular methods (Sinclair et al., 2018; Tsutsui et al., 2020). Conversely, treating half of the root system with deficient medium allows detecting a shoot deficiency response while still sufficiently supplied by the other half of the root, and thus characterizing a root-to-shoot deficiency signal. In the case of a long-distance signal triggered upon resupply of deficient plants, it is delicate to distinguish signaling from delayed nutrient flux in such experimental setups and alternative approaches will need to be designed to identify the putative signal.

In summary, our study revealed the complexity of the zinc homeostasis network in Brachypodium by comparing static and dynamic responses to zinc supply. We identified a short-lived zinc shock response to zinc resupply in roots and hypothetical long-distance zinc signaling that could be important in realistic field resupply conditions. The study also showed that Brachypodium responds phenotypically and genetically to changes in zinc supply, and represents a valuable model of staple grass crops to examine zinc homeostasis that contrasts with the widely studied model Arabidopsis. Differences in zinc/iron interactions and in dynamics of transcriptional changes upon zinc resupply reveal the diversity of zinc homeostasis mechanisms among plant species.

## Supporting information

Amini et al. Supplemental Data

## Acknowledgements

Funding was provided by F.R.S.-FNRS grants MIS F.4511.16, CDR J.0009.17 and PDR T0120.18 (MH). Marie Schloesser, Marie Scheuren, Cédric Delforge, Ana Galinski, Berndt Kastenholz, and Tanja Ehrlich are thanked for technical assistance. The GIGA Genomics platform (ULiege) and the Durandal cluster (InBioS-PhytoSystems, ULiege) are thanked for access to sequencing and computational resources, respectively. SA is funded by a PhD grant of FZJ. MH is a Senior Research Associate of F.R.S.-FNRS. No conflict of interest.

## Author contributions

MH, SA and BA designed the research. SA performed experiments. SA, MH, BA analyzed the data. SG, MC, BB and PM contributed to data interpretation. SA made the figures. SA, MH and BA wrote the manuscript with contributions by MW. All authors read and approved the manuscript.

## Notes

### Competing Interest Statement

The authors have declared no competing interest.

