## Supplementary material for "Transcriptional regulation of *ZIP* genes is independent of local zinc status in Brachypodium shoots upon zinc deficiency and resupply": Amini et al. Supplemental Data

**Supplementary Datasets, Table and Figures**

The following datasets are provided as additional Excel sheets.

**Data S1.** Normalized read counts of all genes in RNA sequencing

**Data S2.** Root and shoot list of DEG in 9 contrasts

**Data S3.** Root and shoot GO enrichment data

**Data S4.** Root and shoot gene clusters

**Data S5.** Root and shoot ZIP cluster, signaling BPs enriched in 10 min in root

**Data S6.** ICP-MS data

**Data S7.** Transcription factor-related genes among DEG lists

**Table S1**. Primers designed and used for quantitative RT-PCR (sequences are 5' to 3')

| Gene | Primer name | Primer sequence | Primer efficiency |
|---|---|---|---|
| Bradi1g12860 | BdIRT1_F | TCCACCAGATGTTCGAAGGC | 1.703 |
| | BdIRT1_R | TCATCTTGGTGCCGTACTCG | |
| Bradi1g17090 | BdNAS_F | GTGCAGAAGATAACCAAGCTCG | 1.892 |
| | BdNAS_R | AAGAGCGAGTTGACTTCCGG | |
| Bradi1g33347 | BdHMA1_F | TGTGGCCTTGTCATTGGTTG | 1.962 |
| | BdHMA1_R | CCCCTACAGACTGAGTTACCGA | |
| Bradi1g53680 | BdZIP13_F | TCGCGTAATCTCTCAGGTCC | 1.998 |
| | BdZIP13_R | TTGATGGTGTCCGGTTCCTG | |
| Bradi2g33110 | BdZIP7_F | GGGGATGTATCAGAGCACGT | 1.851 |
| | BdZIP7_R | ATTCCCATCTCCAGTATCTGTG | |
| Bradi3g17900 | BdZIP4_F | CCGCAGATTTCAACAACCCG | 1.977 |
| | BdZIP4_R | CGAGGTATGTTGCGAGCTGA | |
| Bradi4g00660 | BdUBC18_F | GGAGGCACCTCAGGTCATTT | 1.936 |
| | BdUBC18_R | ATAGCGGTCATTGTCTTGCG | |
| Bradi1g06860 | BdEF1a_F | CCATCGATATTGCCTTGTGG | 1.988 |
| | BdEF1a_R | GTCTGGCCATCCTTGGAGAT | |

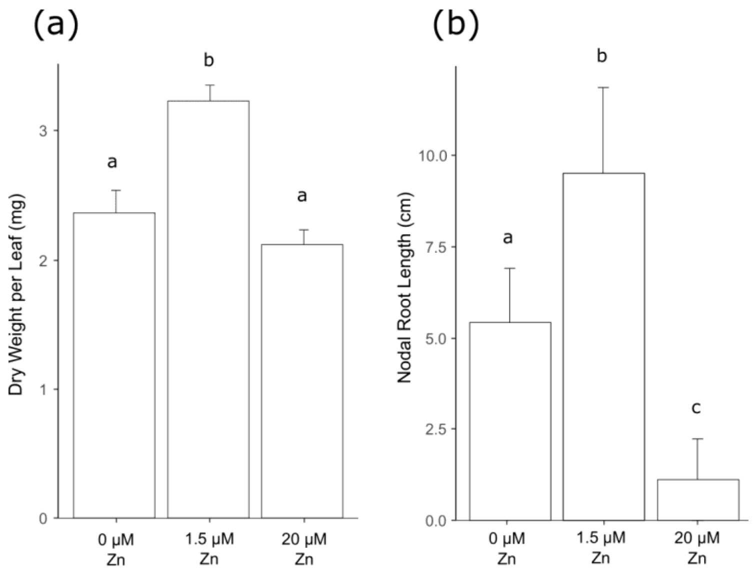


**Figure S1.** Additional shoot and root phenotypic measures of Brachypodium plants upon zinc deficiency and excess. Plants grown hydroponically were exposed for 3 weeks to zinc deficiency (0 µM Zn), control (1.5 µM Zn) or excess (20 µM Zn) conditions. (a) Dry weight per leaf, and (b) Nodal root length. Bars show mean values (+/- standard deviation) of 9-12 individual plants. Letters indicate statistical differences (*p*-value < 0.05) according to Student's T-test.

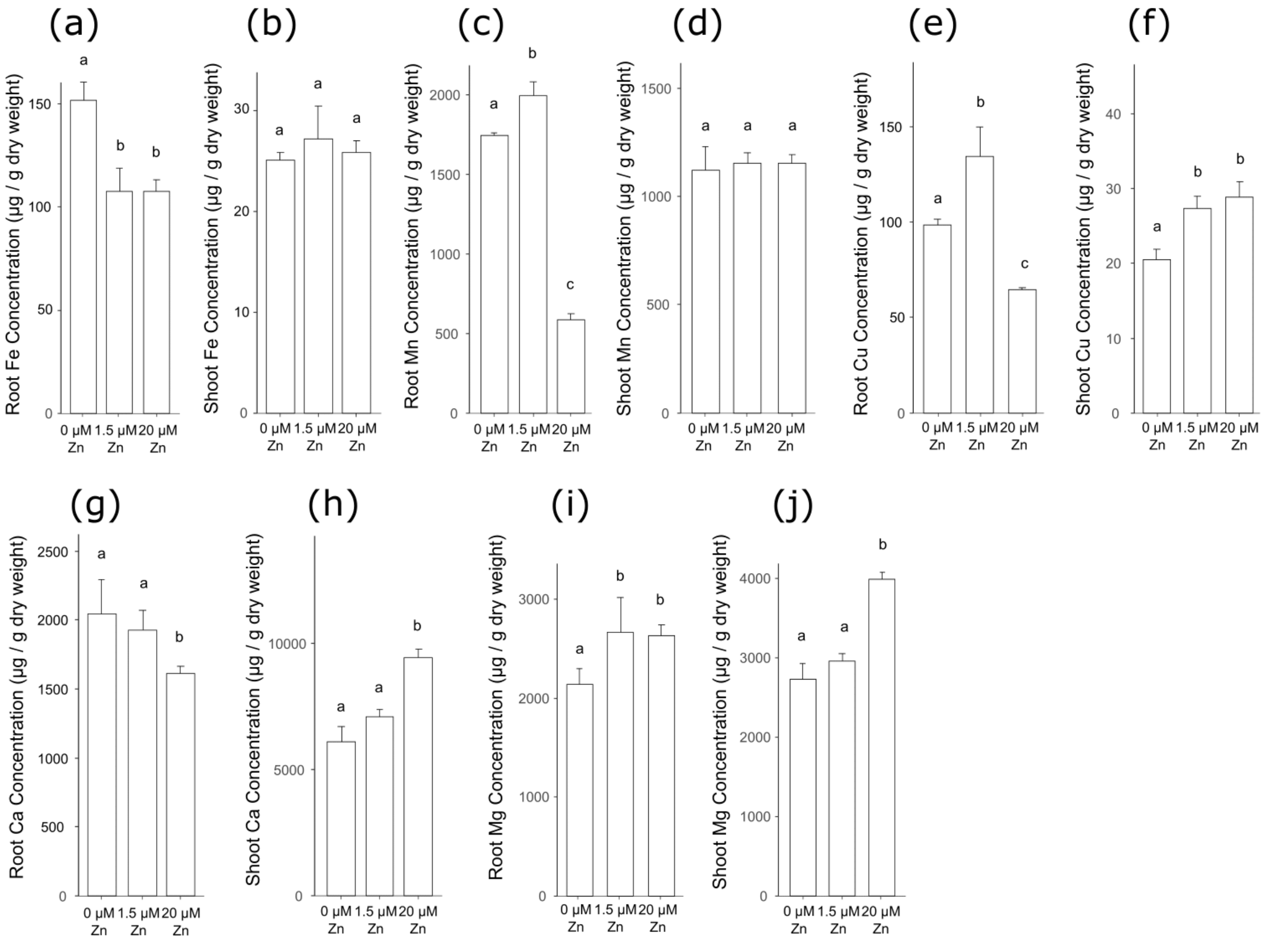


**Figure S2.** Ionome profiling of roots and shoots of Brachypodium upon zinc deficiency and excess. (a) Root iron (Fe), (b) shoot iron, (c) root manganese (Mn), (d) shoot manganese, (e) root copper (Cu), (f) shoot copper, (g) root calcium (Ca), (h) shoot calcium, (i) root magnesium (Mg), and (j) shoot magnesium concentrations. Bars show mean values (+/- standard deviation) of three biological replicates (3-4 plants each). Letters indicate statistical differences ($p$-value < 0.05) according to Student's T-test.

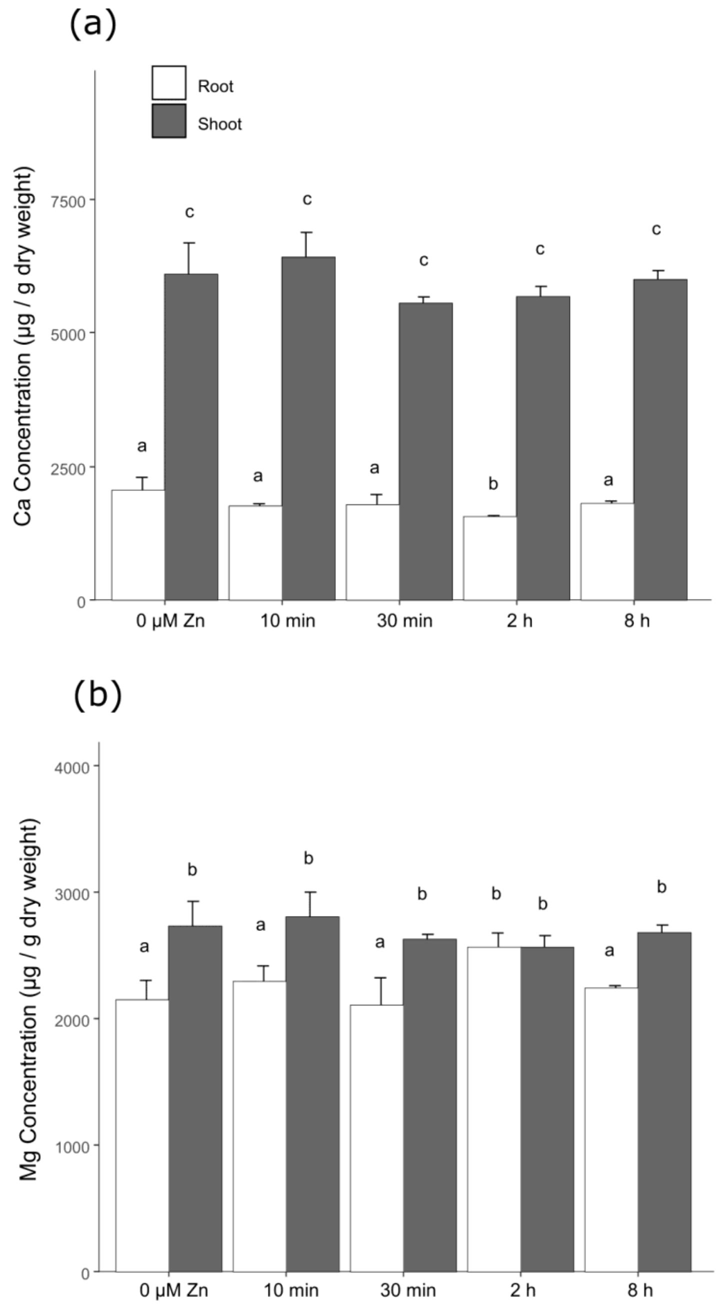


**Figure S3.** Profiling of calcium and magnesium accumulation in roots and shoots of Brachypodium upon zinc deficiency and re-supply. Plants grown hydroponically under zinc deficiency (0 µM Zn) for 3 weeks were resupplied with 1 µM Zn and samples were harvested after short time points (10 minutes to 8 hours). Root and shoot (a) calcium (Ca), and (b) magnesium (Mg) concentration of plants. Bars show mean values (+/- standard deviation) of three biological replicates (3-4 plants each). Letters indicate statistical differences ($p$-value < 0.05) according to one-way ANOVA.

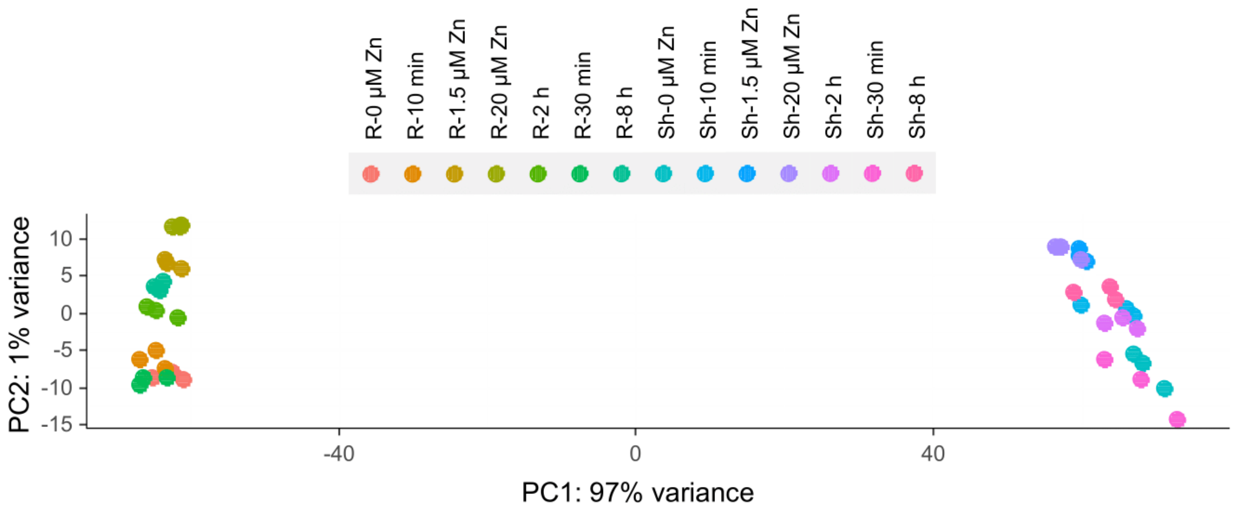


**Figure S4.** Principal Component Analysis of combined root and shoot RNA sequencing expression data. RNA sequencing analysis was used to examine the steady-state response to zinc deficiency and excess and the dynamic response to zinc deficiency and resupply in Brachypodium. “R”: root samples, “Sh”: shoot samples.

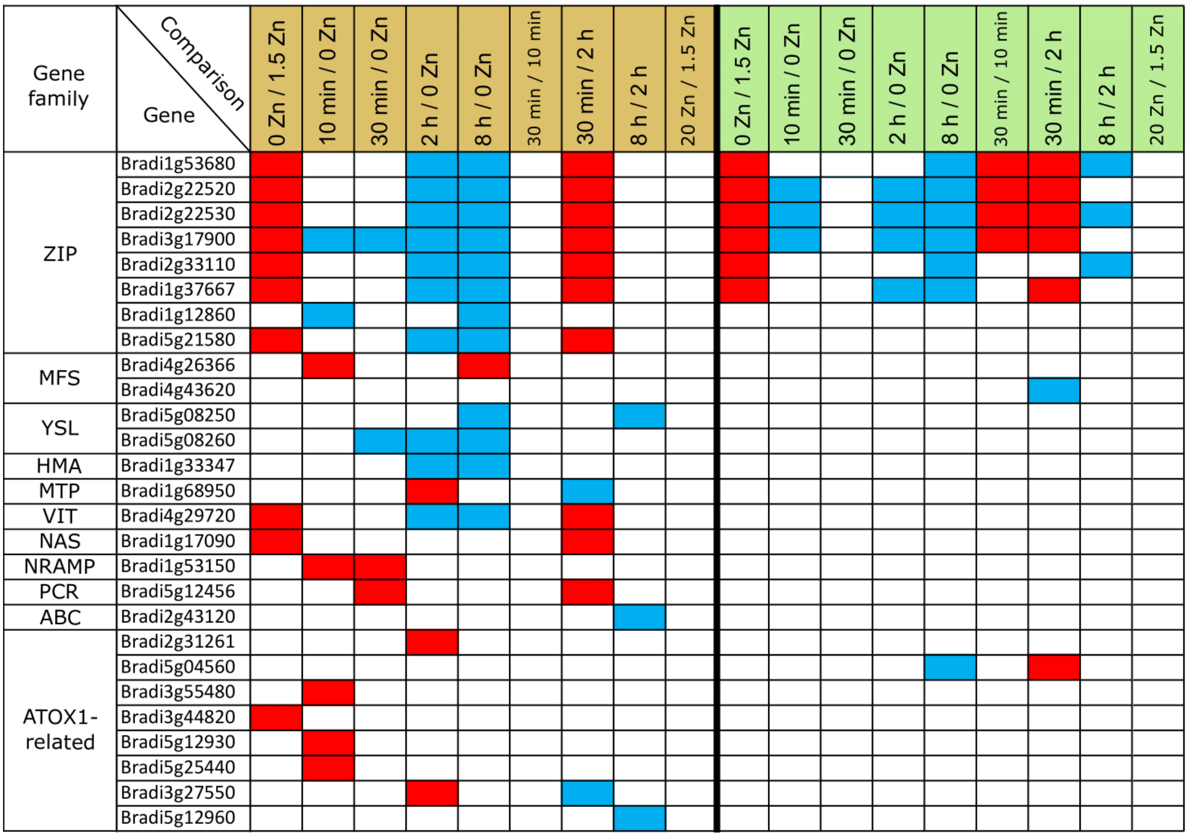


**Figure S5.** Map of differentially expressed metal-related genes in the steady-state response to zinc deficiency and excess and the dynamic response to zinc deficiency and resupply in Brachypodium. The data are presented for 9 selected comparisons. In the header, brown and green colors show root and shoot samples, respectively. Up- and down-regulated genes are in red and blue, respectively.

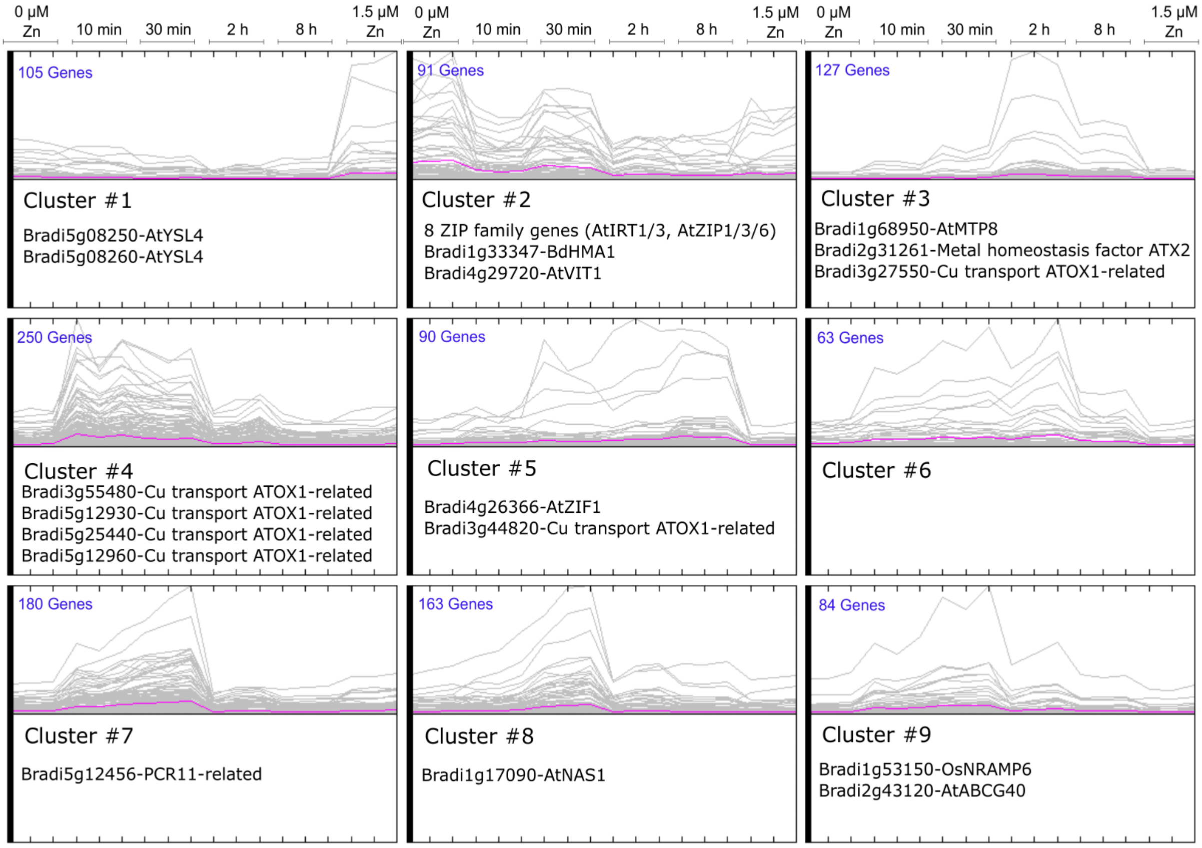


**Figure S6.** Clustering of gene expression in roots upon zinc deficiency and resupply in Brachypodium. Pearson correlation was used as distance metric in k-means clustering. The number of differentially expressed genes included in each cluster is noted in blue in each box. Lines are there to indicate the expression profile of the genes across the three biological replicates, and they should not be considered as time progression. The pink lines show the mean expression of all genes in each cluster. Metal homeostasis-related genes present in the clusters are noted under each graph. Cluster #2 included in Fig. 8 is presented again for completeness.

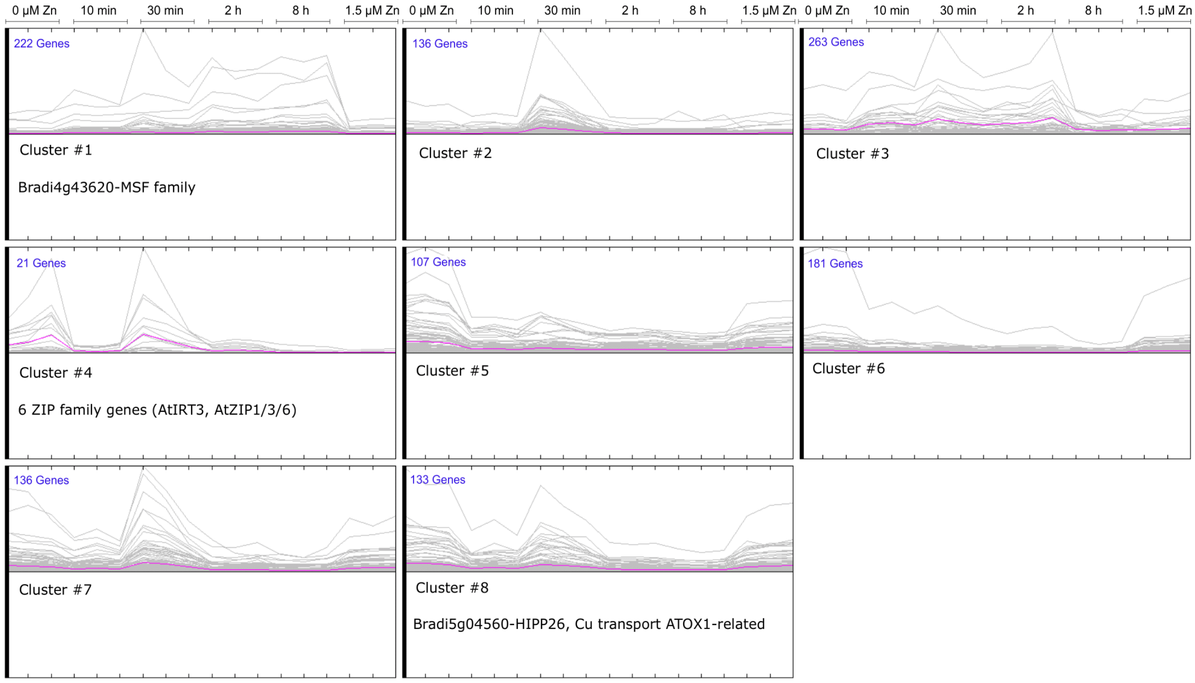


**Figure S7.** Clustering of gene expression in shoots upon zinc deficiency and resupply in Brachypodium. Pearson correlation was used as distance metric in k-means clustering. The number of differentially expressed genes included in each cluster is noted in blue in each box. Lines are there to indicate the expression profile of the genes across the three biological replicates, and they should not be considered as time progression. The pink lines show the mean expression of all genes in each cluster. Metal homeostasis-related genes present in the clusters are noted under each graph. Cluster #4 included in Fig. 8 is presented again for completeness.

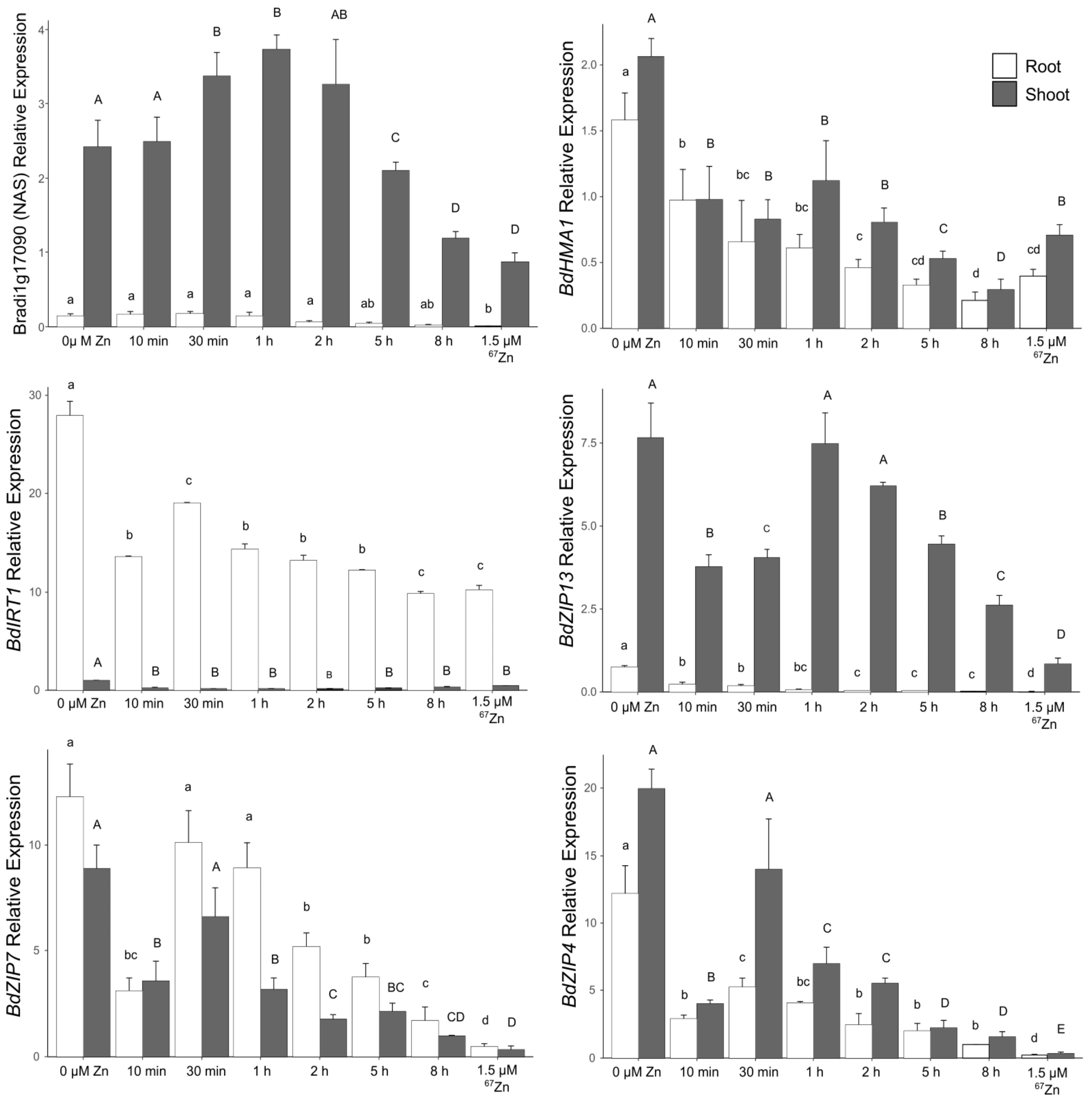


**Figure S8.** Relative gene expression of metal homeostasis genes upon zinc deficiency and $^{67}$Zn resupply in Brachypodium. Root and shoot transcript levels of the Bradi1g17090 (NAS family), *BdHMA1*, *BdIRT1*, *BdZIP13*, *BdZIP7* and *BdZIP4* genes were determined by quantitative RT-PCR. Expression levels are relative to *UBC18* and *EF1α*, and scaled to average. Bars show mean values (+/- standard deviation) of three biological replicates (2-3 plants each). Letters indicate statistical differences ($p$-value < 0.05) according to one-way ANOVA.

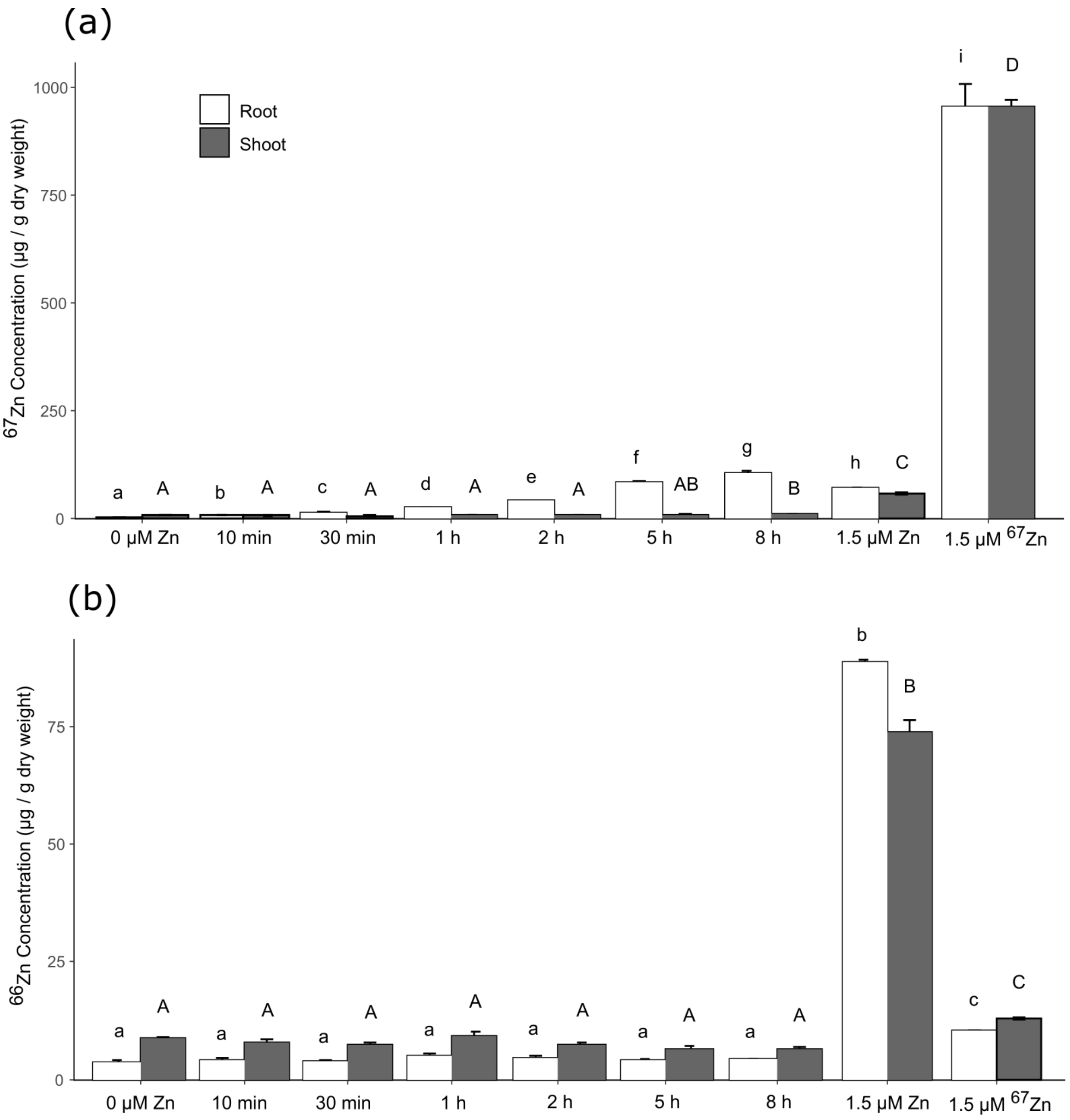


**Figure S9.** Elemental analysis of zinc isotopes in Brachypodium plants throughout the time series upon $^{67}$Zn resupply. (a) Root and shoot $^{67}$Zn, and (b) $^{66}$Zn concentrations. $^{66}$Zn concentration was measured as a control. Supplying plants with natural zinc (Zn) was another control for this experiment. Bars show mean values (+/- standard deviation) of three biological replicates (4 plants each). Letters indicate statistical differences ($p$-value < 0.05) according to one-way ANOVA.

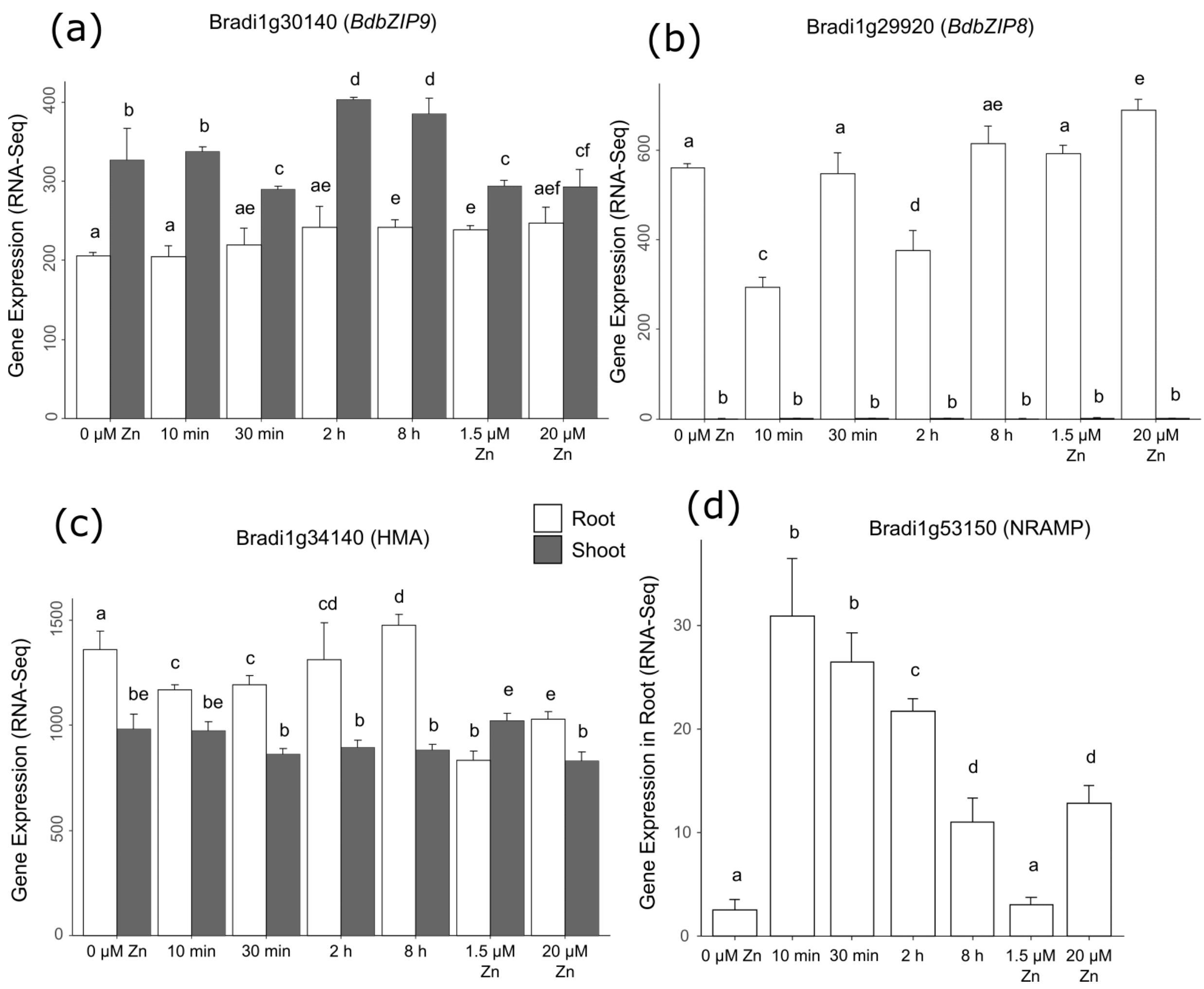


**Figure S10.** Selected candidate gene expression levels upon zinc deficiency and resupply in Brachypodium. Root and shoot (a) Bradi1g30140 (*BdbZIP9*), and (b) Bradi1g29920 (*BdbZIP8*), and (c) Bradi1g34140 (HMA) transcript levels. (d) Root transcript level of Bradi1g53150 (NRAMP). Bars show mean values (+/- standard deviation) of three biological replicates (3-4 plants each) from RNA Sequencing data. Letters indicate statistical differences ($p$-value < 0.05) according to one-way ANOVA.
